# A shift to shorter cuticular hydrocarbons accompanies sexual isolation among *Drosophila americana* group populations

**DOI:** 10.1101/2020.09.07.286294

**Authors:** Jeremy S. Davis, Matthew J. Pearcy, Joanne Y. Yew, Leonie C. Moyle

**Author notes:** RRH: Sexual isolation and chemical signal evolution.

## Abstract

Because sensory signals often evolve rapidly, they could be instrumental in the emergence of reproductive isolation between species. However, pinpointing their specific contribution to isolating barriers, and the mechanisms underlying their divergence, remains challenging. Here we demonstrate sexual isolation due to divergence in chemical signals between two populations of *Drosophila americana* (SC and NE) and one population of *D. novamexicana*, and dissect its underlying phenotypic and genetic mechanisms. Mating trials revealed strong sexual isolation between *Drosophila novamexicana* males and SC *Drosophila americana* females, as well as more moderate bi-directional isolation between *D. americana* populations. Mating behavior data indicates SC *D. americana* males have the highest courtship efficiency and, unlike males of the other populations, are accepted by females of all species. Quantification of cuticular hydrocarbon (CHC) profiles—chemosensory signals that are used for species recognition and mate finding in *Drosophila*—shows that the SC *D. americana* population differs from the other populations primarily on the basis of compound carbon chain-length. Moreover, manipulation of male CHC composition via heterospecific perfuming—specifically perfuming *D. novamexicana* males with SC *D. americana* males—abolishes their sexual isolation from these *D. americana* females. Of a set of candidates, a single gene—elongase CG17821—had patterns of gene expression consistent with a role in CHC differences between species. Sequence comparisons indicate D. novamexicana and our Nebraska (NE) *D. americana* population share a derived CG17821 truncation mutation that could also contribute to their shared “short” CHC phenotype. Together, these data suggest an evolutionary model for the origin and spread of this allele and its consequences for CHC divergence and sexual isolation in this group.

## Introduction

Sensory signals can act as sexual cues that are critical for intraspecific mate evaluation and reproductive success. Moreover, because they are frequently among the most rapidly evolving species differences (Smadja and Butlin 2009, Seddon et al 2013, Wilkins et al 2013), divergence in sensory sexual signals might often contribute to the earliest stages of reproductive isolation, in the form of prezygotic barriers between lineages (Butlin et al 2012, Ritchie 2007). Among all such sexual signals, nowhere is the diversity more evident than those acting as premating traits, including coloration and other visual signals, acoustic signals, and complex pheromone blends. However, convincingly demonstrating the connection between such sensory signal divergence and emerging reproductive isolation can be challenging because it requires, first, identification and demonstration of the direct role of specific signals mediating sexual isolation between species and, second, understanding the specific mechanistic changes that have given rise to lineage differences in this signal.

Among insects, sexual signals draw on three primary sensory modalities—visual, auditory, and chemosensory—and most species likely use a combination of all three modalities to identify potential mates. Of these, many insect chemosensory signals primarily consist of cuticular hydrocarbons (CHCs)— a broad group of carbon-chain compounds important for many essential functions including sex pheromone signaling, but also environmental adaptation (Blomquist and Bagneres 2010, Chung and Carrol 2015, Yew and Chung 2017). Unlike many auditory and visual cues, CHCs are produced and received by both males and females, making them potentially important for both male and female mate choice or species recognition. Indeed, studies on *Drosophila* chemical communication, particularly in the melanogaster subgroup—have detected clear evidence for sexual isolation based on male choice of female CHCs (Coyne et al. 1995, Billeter et al. 2009), while work in other species has demonstrated female choice of species-specific male CHC profiles (Coyne et al. 2002, Mas and Jallon 2005, Curtis et al. 2013, Dyer et al. 2014). Elements of the biochemical pathways of CHC production and the underlying genes are also relatively well understood in D. melanogaster (Pardy et al 2018). Accordingly, single genes contributing to species-specific CHC differences have been implicated as causes of reproductive isolation in several cases, including between D. melanogaster and *D. simulans* (desatF: Legendre et al. 2008) and *D. serrata* and *D. birchii* (mFAS: Chung et al. 2014). This framework provides an excellent resource for identifying candidate genes for sensory sexual signal variation in other *Drosophila* systems.

Here we use the *Drosophila americana* group to investigate how species variation in CHC profiles contributes to reproductive isolation, and to identify the underlying chemical and genetic changes that may comprise this variation. This group includes two closely related (MRCA ∼0.5 mya, Morales-Hojas et al. 2008) species—*D. novamexicana* and *D. americana*—that occupy distinct geographic and environmental habitats (Davis and Moyle 2019). *D. americana* is broadly distributed in the United States from the east coast to the Rocky Mountains, and exhibits significant phenotypic and genetic variation among populations throughout (e.g., Caletka and McAllister 2004, Davis and Moyle 2019), while *D. novamexicana* is localized to the arid southwestern US. Both species have been noted as being associated with—and exclusively collected near—willow species of the genus *Salix*, though the exact nature of this association is not known (Blight and Romano 1953, McAllister 2002). Classical mating studies (Spieth 1951) have documented population-specific variation in reproductive isolation between members of this group (see supplemental Table S1 and below). Prior analysis has also shown qualitative differences in CHC composition between males (Bartelt at al. 1986), however the contribution of hydrocarbons to premating isolation has not been assessed. With three populations from this group, one *D. novamexicana* population and two *D. americana* populations—one southern (South Carolina, hereafter SC) and one western (Nebraska, hereafter NE)—here we use a combination of mating and behavioral studies, chemical analysis and manipulation, and gene expression and sequence variation analyses to assess the role of CHCs in reproductive isolation. Together, our data suggest that evolution of a male sexual signal—an overall shift in the relative abundance of longer versus shorter cuticular hydrocarbons, due to novel mutation in an elongase gene—has produced complete premating isolation between derived males and females from species that retain the ancestral trait and preference, as proposed in classical models (Kaneshiro 1976, 1980) of the evolution of asymmetric sexual isolation.

**Table 1:** Copulation rates and latency for each cross in unperfumed 4x4 and 1x1 mating trials. Isolation index included is RI_1_ as defined by Sobel and Chen (2014), and uses values from the 4x4 mating trials.

| Female species | Male species | 4x4 trials | 1x1 trials | | Isolation Index ($RI_1$) |
| --- | --- | --- | --- | --- | --- |
| | | Mating rate<sup>a</sup><br>(mean ± SE) | Mating rate<sup>a</sup><br>(mean) | Cop. latency<br>(mean ± SE)<sup>b</sup> | |
| *D. novamexicana* | *D. novamexicana* | 0.90 ± 0.06 | 1.00 | 69.89 ± 22.73 | |
| | NE *D. americana* | 0.69 ± 0.12 | 0.60 | 147.48 ± 27.00 | 0.233 |
| | SC *D. americana* | 0.71 ± 0.18 | 0.60 | 83.63 ± 39.70 | 0.211 |
| NE *D. americana* | *D. novamexicana* | 1.00 ± 0 | 0.20 | 158.33 ± 21.67 | 0 |
| | NE *D. americana* | 1.00 ± 0 | 0.60 | 123.46 ± 29.01 | |
| | SC *D. americana* | 0.50 ± 0.18 | 0.60 | 99.42 ± 34.74 | 0.5 |
| SC *D. americana* | *D. novamexicana* | 0.00 ± 0 | 0.00 | 180 ± 0 | 1 |
| | NE *D. americana* | 0.40 ± 0.14 | 0.60 | 99.42 ± 34.74 | 0.52 |
| | SC *D. americana* | 0.85 ± 0.06 | 0.80 | 50.32 ± 33.07 | |

<sup>a</sup> n = 5, rate is a proportion out of maximum possible matings (4 or 1)
<sup>b</sup> n = 5, units are in minutes. Unmated trials are scored as a latency of 180 min—the maximum time observed.

## Results

### SC *D. americana* females discriminate against heterospecific males in mating trials

We found clear evidence of moderate to strong sexual isolation between SC *D. americana* and the other two populations (Table 1). Mating rate (average proportion of females mated, in 4x4 mating trials) ranged from very high in intrapopulation crosses and some interpopulation crosses, to <50% in interpopulation combinations of female SC *D. americana* with each of the other species. Notably, *D. novamexicana* males were never successful in mating with SC *D. americana* females, indicating strong sexual isolation in this cross direction between these populations. In contrast, mating rate in the reciprocal cross was 70%. The other pairing with mating rates at or below 50% involved both reciprocal directions between *D. americana* populations. Across all cross combinations in the 4x4 mating trials, we found that female identity (Kruskal-Wallis test: χ2 (2) = 8.99, *P* = 0.012), but not male identity (Kruskal-Wallis test: χ2 (2) = 0.80, *P* = 0.67), significantly affected success. Post-hoc comparisons confirmed no pairwise differences in male mating rate between any populations (Table S2). In contrast, the copulation rate of SC *D. americana* females was significantly lower on average than both NE *D. americana* (P-adj = 0.025) and *D. novamexicana* (P-adj = 0.042), whereas NE *D. americana* and *D. novamexicana* females did not differ (P-adj = 0.15). Together, these results indicate that SC *D. americana* females discriminate against heteropopulation males more strongly than do females of the other two populations, with the greatest discrimination against *D. novamexicana* males. This discrimination produces strongly asymmetric sexual isolation between SC *D. americana* and *D. novamexicana*, while premating isolation between the two *D. americana* populations is bi-directional and more moderate. Importantly, this strong asymmetric isolation reiterates results documented by Spieth (1951, Table S1), using similar 4x4 mating experiments with 2 *D. novamexicana* and 9 *D. americana* lines—including the NE *D. americana* and *D. novamexicana* populations used here. In this prior analysis, males from *D. novamexicana* showed little to no mating success with southeastern *D. americana* populations (previously referred to as D. a. texana) (0-12.5% mating frequency; Table S1), while *D. novamexicana* females were more receptive in the reciprocal cross (mating frequency of 46.7-86.5%; Table S1). The consistency of these results 70 years apart suggests isolation observed here is not a bi-product of long-term laboratory culture, but instead reflects differences present in the natural populations from which these lines were collected.

**Table 2:** Mean copulation success (number of matings out of 4 per trial) with SC *D. americana* females for each perfuming identity in manipulated (perfumed) 4x4 mating assays

| Target male | Donor male | Copulation success<br>(Mean ± SE) | Pairwise Mann-Whitney U test | |
| --- | --- | --- | --- | --- |
| Nov-SC pair |  |  | U-score | P-value |
| *D. novamexicana* | *D. novamexicana* | 0.25 ± 0.15 | } 0 | 0.0247 |
| *D. novamexicana* | SC *D. americana* | 0.94 ± 0.063 | | |
| SC *D. americana* | SC *D. americana* | 1.00 ± 0 | } 0 | 0.018 |
| SC *D. americana* | *D. novamexicana* | 0.19 ± 0.063 | | |
| NE-SC pair |  |  |  |  |
| NE *D. americana* | NE *D. americana* | 0.44 ± 0.12 | } 5.5 | 0.54 |
| NE *D. americana* | SC *D. americana* | 0.56 ± 0.12 | | |
| SC *D. americana* | SC *D. americana* | 1.00 ± 0 | } 0 | 0.018 |
| SC *D. americana* | NE *D. americana* | 0.50 ± 0.063 | | |

Note: n = 5 for all crosses

### SC *D. americana* males have greater courtship efficiency and mating success

We video-recorded single-pair matings for each cross combination to evaluate population differences in courtship strategies and whether these differed in interpopulation pairings. Three male courtship behaviors were quantified from these 1x1 trials—display rate, tapping rate, and licking rate. All three behaviors showed similar patterns of among-population variation, when each was evaluated for the effect of male population, female population, or their interaction, using 2-way ANOVAs. Males significantly differed with respect to display-rate (*F*(2, 36) = 3.76, *P* = 0.03) and tap-rate (*F*(2, 36) = 4.76, *P* = 0.017), but only marginally for lick-rate (*F*(2, 36) = 2.67, *P* = 0.083). For all three behaviors, this difference is due to SC *D. americana* males exhibiting higher rates (Tukey HSD post-hoc tests; Figure 1, supplementary Table S3). In contrast, we detected no female identity effects on male behavioral rate (display-rate: F(2, 42) = 1.83, *P* = 0.18; tap-rate F(2, 36) = 0.58, *P* = 0. 56; lick-rate: F(2, 36) = 0.43, *P* = 0.65), or any female x male interactions (display-rate: F(2, 36) = 0.72, *P* = 0.58; tap-rate F(2, 36) = 0.45, *P* = 0.77; lick-rate: F(2, 36) = 0.65, *P* = 0.63). Because behavior rates are dependent on copulation latency, and not simply the total number of behavioral events, these rates approximate the efficiency with which each courtship behavior results in a mating. Taken together, our results indicate that SC *D. americana* males have greater courtship efficiency (in terms of display and tap rates), and that males do not significantly vary these behavioral rates depending on female population identity.

**Figure 1:**
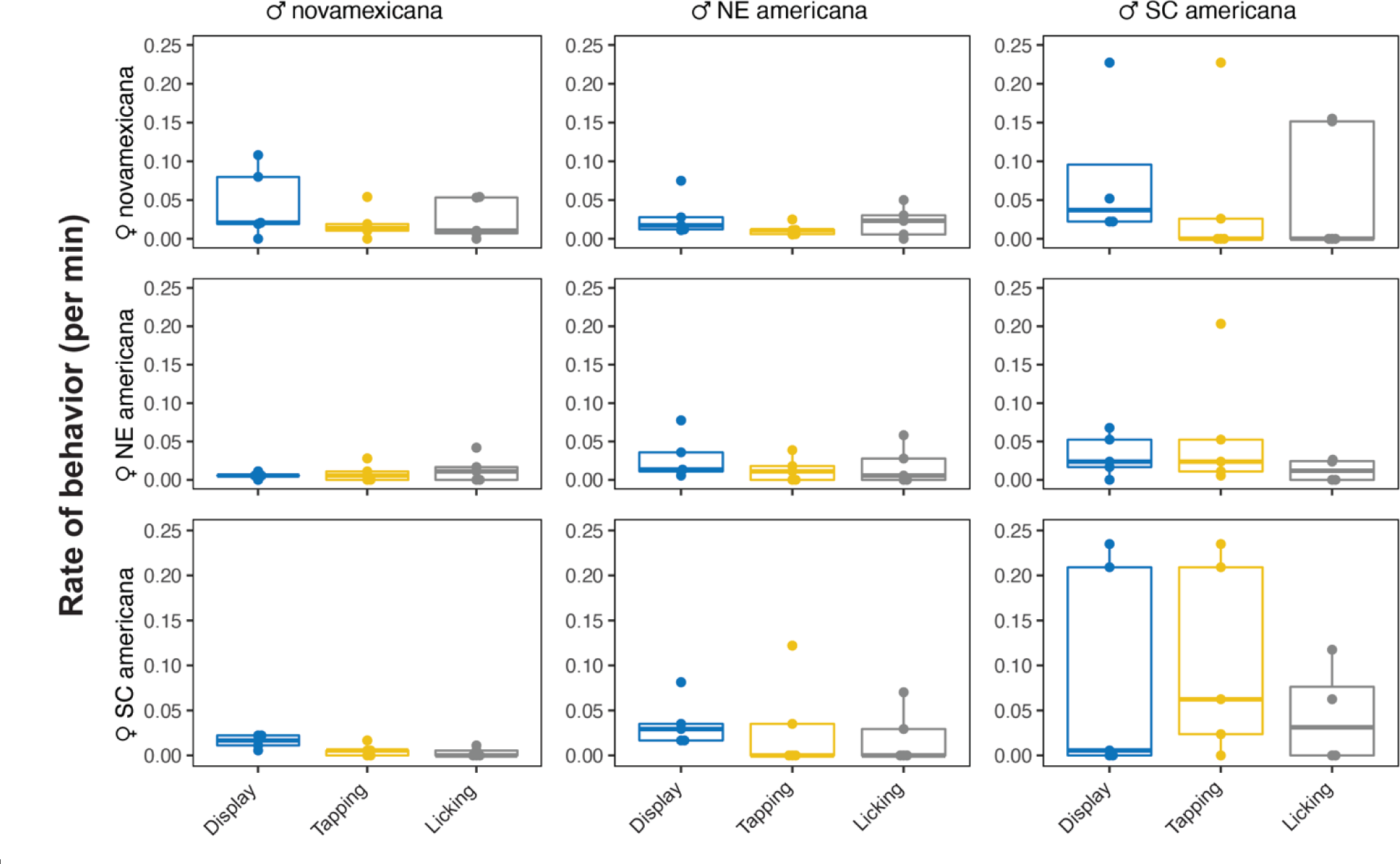
Male courtship behavior rates in 1x1 mating assays across all cross types. Points represent individual trials, with boxes showing quartiles and the mean as a solid bar. Males (columns) show significant differences in display rate and tap rate, and marginal differences in lick rate. Rates did not differ based on female identity (rows) or male x female interaction (see results). Bar and whiskers indicate mean and standard error (SE).

**Table 3:** Molecular evolution (*d*_N_/*d*_S_) and species differences in quantitative gene expression for 23 candidate genes associated with CHC function, analyzed separately by sex, with *F*-value of ANOVA test and rank compared to all genes in dataset. *d*_N_/*d*_S_ of all genes reported in supplementary file

| Candidate gene | *D. virilis* ortholog | InterProt domain | $d_N/d_S^a$ | Males | | Females | |
| --- | --- | --- | --- | --- | --- | --- | --- |
| | | | | $F$-value<sup>b</sup> | Rank %ile<sup>b</sup> | $F$-value<sup>b</sup> | Rank %ile<sup>b</sup> |
| Cyp4g1 | GJ15981 | Cyp4g | 0.0469 | 0.143 | 95% | 0.158 | 87% |
| Cyp4g15 | GJ19152 | Cyp4g | 0.0862 | 2.07 | 33% | 1.05 | 38% |
| Cyt-b5-r | Cyt-b5-r | fatty-acid desaturase | 0.0001 | 2.32 | 29% | 0.736 | 53% |
| ifc | GJ15437 | fatty-acid desaturase | 0.0266 | 0.122 | 96% | 0.74 | 52% |
| CG17928 | GJ17306 | fatty-acid desaturase | 0.133 | 1.09 | 56% | 0.0233 | 96% |
| CG8630 | GJ10413 | fatty-acid desaturase | 0.118 | 3.48 | 19% | 1.46 | 27% |
| CG9743 | GJ10408 | fatty-acid desaturase | 0.0712 | 2.83 | 24% | 1.75 | 22% |
| desat2 | desat2 | fatty-acid desaturase | 0.0780 | 0.216 | 93% | 0.903 | 45% |
| Baldspot | GJ12451 | elongase | 0.0280 | 0.814 | 68% | 0.916 | 45% |
| bond | GJ24166 | elongase | 0.0503 | 0.710 | 53% | 0.949 | 79% |
| **ELOVL** | **GJ24115** | **elongase** | 0.0291 | **9.61** | **6%** | 2.67 | 14% |
| CG5326 | GJ24664 | elongase | 0.0081 | 4.14 | 16% | 0.138 | 89% |
| **CG17821** | **GJ22070** | **elongase** | **0.284** | 3.50 | 19% | **17.2** | **0.8%** |
| **CG18609** | **GJ22071** | **elongase** | **0.249** | **8.19** | **7%** | 1.85 | 21% |
| **CG30008** | **GJ22296** | **elongase** | 0.151 | 2.19 | 31% | **4.33** | **7%** |
| CG31522 | GJ24118 | elongase | 0.0422 | 0.744 | 71% | 0.380 | 73% |
| **CG31523** | **GJ24117** | **elongase** | 0.0410 | **6.57** | **9%** | 0.0481 | 95% |
| CG33110 | GJ24167 | elongase | 0.0178 | 0.466 | 83% | 0.436 | 70% |
| **CG6660** | **GJ14350** | **elongase** | **0.387** | 1.15 | 54% | 1.20 | 33% |
| FASN1 | GJ21736 | fatty-acid-synthase | 0.0274 | 1.42 | 46% | 0.526 | 64% |
| FASN2 | GJ21725 | fatty-acid-synthase | 0.0979 | 1.83 | 37% | 1.02 | 39% |
| ebony | GJ14444 | pigment | 0.0280 | 0.673 | 33% | 1.15 | 35% |
| tan | GJ14875 | pigment | 0.0269 | 1.49 | 45% | 0.347 | 75% |

<sup>a</sup>**Bold** genes in this column indicate $d_N/d_S$ values over 1 standard deviation from mean $d_N/d_S$ value of 4397 genes analyzed.
<sup>b</sup>**Bold** genes in these columns indicate genes in the top 10% of differentially expressed genes within sex.

The single pair mating assays also broadly reiterated the patterns of moderate to strong sexual isolation we observed for SC *D. americana* in 4x4 trials. As with our 4x4 mating assay, *D. novamexicana* males never mated with SC *D. americana* females when paired individually, while mating rate between the two *D. americana* populations was moderately reduced in both directions. A logistic regression assessing how mating rate varied based on male identity, female identity, and their interaction showed a significant interaction effect only (males: χ2 (2) = 2.97, *P* = 0.28; females: χ2 (2) = 2.52, *P* = 0.23; males*females: χ (4) = 12.67, *P* = 0.013)—a difference from 4x4-mating trials where female population identity was the primary predictor of mating rate. This variation between the assays appears to be driven by female NE *D. americana* accepting fewer *D. novamexicana* males in 1x1 trials (20%) compared to 4x4 trials (100%, Table 1), suggesting that male density within mating trials might affect copulation success in this particular species combination. Finally, copulation latency differed marginally based on both male identity (ANOVA: F(2) = 3.14, *P* = 0.055) and the male by female interaction (*F*(4) = 2.48, *P* = 0.062), but did not differ based on female species identity (*F*(2) = 0.68, *P* = 0.5147). Post-hoc tests suggest that the marginal male population effect is likely due to lower copulation latency in SC *D. americana* males relative to *D. novamexicana* males (Table S4).

### Populations and sexes differ in CHC composition

We found that both populations and sexes within populations showed different and distinctive CHC profiles, with the largest differences detected between SC *D. americana* and the other two lines (Figure 2). Across all samples (n=5 replicate samples for each identity) we detected 8 alkene compounds (2 of which are sex-specific) and 4 methyl-branched alkane compounds present in at least one population. A principal component analysis of profiles from unmanipulated flies (unmanipulated-PCA or ‘U-PCA’) summarizing the primary axes of CHC profile composition found that 95.3% of compound variation across all samples was explained by the first three principal components (U-PC1 to 3). Of these, CHC composition varied significantly for both population and sex for U-PC1 (pop: F(2, 429.29), P < 0.0001; sex: F(1, 46.97), P < 0.0001) and U-PC2 (pop: F(2, 10.54), *P* = 0.00043; sex: F(1, 181.69), P < 0.0001), but had no sex or population effect for U-PC3 (pop: F(2, 0.31), *P* = 0.737; sex: F(1, 0.57), *P* = 0.458)(Figure 2). Notably, SC *D. americana* of both sexes differed from the other two populations along the U-PC1 axis (Tukey HSD; Nov-SC: P-adjust < 0.0001; NE-SC: P-adjust < 0.0001; Nov-NE: P-adjust = 0.28), which also explains most of the CHC variation between samples (69.9%). U-PC1 was positively loaded for most of the shorter carbon chain length compounds in both alkene and methyl-branched alkane classes of compounds, and negative values were strongly loaded for the compounds with longer carbon chain length (Table S5); consequently, this axis can be interpreted as a composite of average compound length across the CHC profile as a whole (Figure 2). Accordingly, both sexes of SC *D. americana* had a higher abundance of longer-chain alkenes and methyl-branched alkanes than did either NE *D. americana* or *D. novamexicana*.

**Figure 2:**
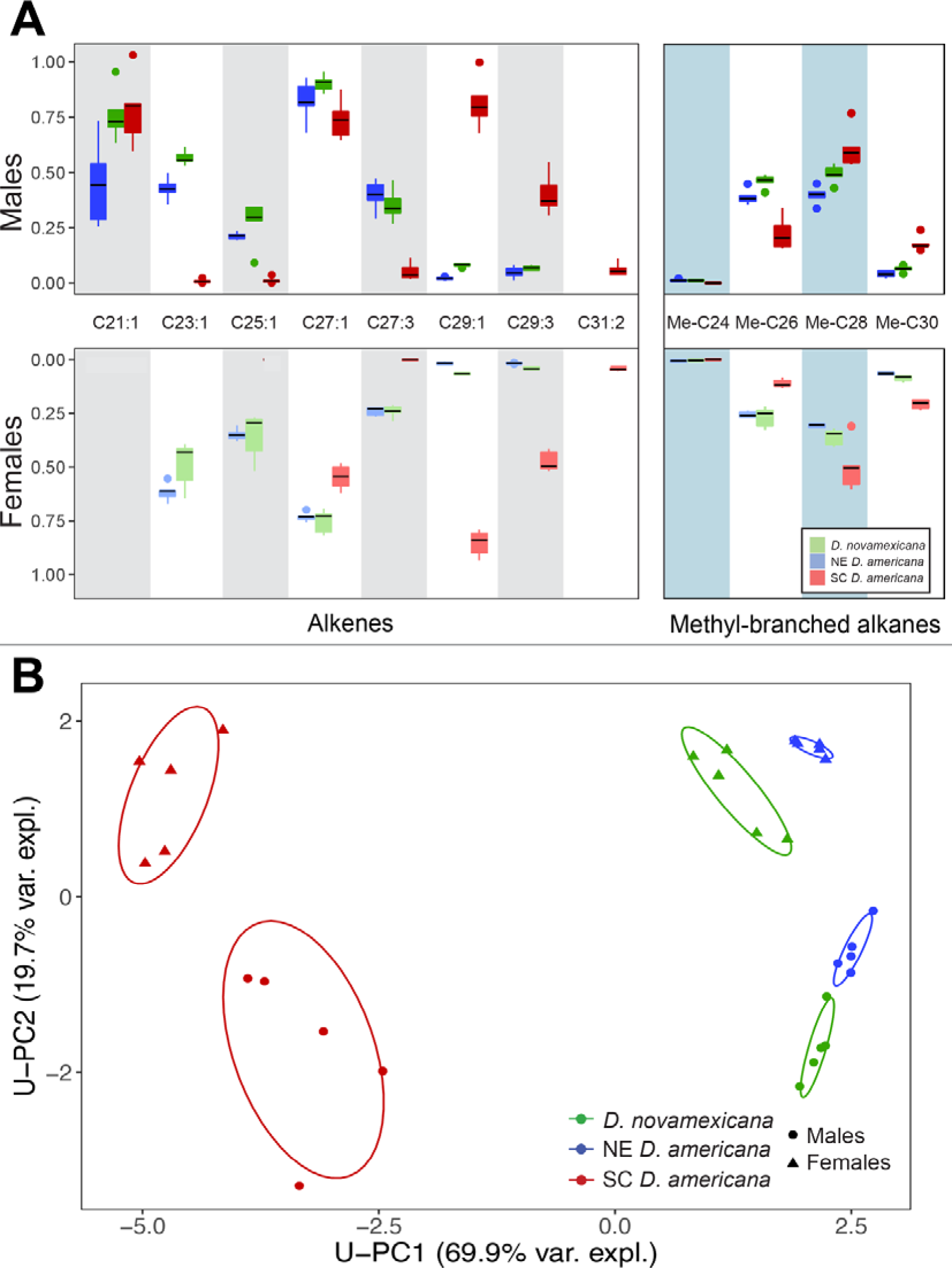
Cuticular hydrocarbon composition of unmanipulated males and females of each species stock. A) Relative log-scale abundance of compounds for males (upper) and females (lower), for each major compound type (line = mean, box = central quartile, whiskers = S.E.). *n* = 5 samples for each sex and line. Test statistics for species differences are in Table S3. B) First two principal components (U-PC1 and U-PC2) of composite CHC variation among males (circle) and females (triangle), with percent of total variance explained. Ellipses indicate 90% bivariate normal density for each species-sex group. Species and sex significantly influenced both U-PC1 and U-PC2.

In contrast to U-PC1, U-PC2 appeared to primarily differentiate sexes within populations, but also NE *D. americana* males from males of the other two populations. This axis was most heavily loaded for two compounds: C21:1 and Me-C28, followed by C27:1 and Me-C26, with much smaller loadings for all other compounds (Table S5). C21:1 is a male-specific compound that is not detected in females; Me-C28, Me-26, and C27:1 abundance was also consistently different between sexes, with males always having more of these CHCs than females within each population (Figure 2). Therefore, negative U-PC2 values can be interpreted as primarily representing more male-specific profiles, while positive U-PC2 values correspond to more female-like profiles. With respect to the detected species difference in U-PC2, post-hoc tests reveal that this was driven by NE *D. americana* (Tukey HSD; Nov-SC: P-adjust = 0.99; NE-SC: P-adjust = 0.0062; Nov-NE: P-adjust = 0.0088). Furthermore, we find that abundance of the individual male-specific compound C21:1 differs between NE *D. americana* and males of the other two populations, but not between *D. novamexicana* and SC *D. americana* (Tukey HSD; Nov-SC: P-adjust = 0.98; NE-SC: P-adjust = 0.016; Nov-NE: P-adjust = 0.022). (Results from tests of individual compound differences can be found in Table S6).

### Interpopulation perfuming influences SC *D. americana* female acceptance of intra- and inter-population males

We found that patterns of sexual isolation could be modified by specifically changing the CHC profiles of males of different populations. To do this, we used perfuming assays to manipulate the pheromone profile of males by co-housing them with either intra- or inter-population males, and then evaluated how this manipulation influenced mating rates among populations. For these analyses, we focused on two pairings in two separate, analogous experiments—*D. novamexicana* and SC *D. americana* (“Nov-SC pair”), and the *D. americana* population pair (“NE-SC pair”)— because SC *D. americana* shows the strongest sexual isolation from the two other populations, and the largest differences in CHC composition. Within each perfuming experiment, we generated four combinations of target and donor male identities: each population perfumed by same-population males (control) and each population perfumed by hetero-population males. Each class of perfumed males was evaluated for mating rate specifically with SC *D. americana* females, because these females were the most discriminating against heteropopulation males in our previous mating assays (Table 2). The observed mating rate of males perfumed with same-population males (the control manipulation) was similar to that observed in unperfumed mating trials: *D. novamexicana* and NE *D. americana* males perfumed with their own population performed poorly with SC *D. americana* females, relative to SC *D. americana* perfumed with their own males, which had a 100% mating rate (Table 2). In strong contrast, hetero-population perfumed males showed altered mating rates relative to control perfumed males. For Nov-SC pairings, *D. novamexicana* males perfumed with SC *D. americana* males successfully mated with SC *D. americana* females 94% of the time, a significantly higher mating rate than *D. novamexicana* males perfumed with their own males (25%; Mann-Whitney U-test: U = 0, *P* = 0.025). (Note that copulation success between conspecific perfumed *D. novamexicana* males and SC *D. americana* remains low (25%) but differs from the complete mating isolation observed in unperfumed trials, possibly because of changes in behavior due to male-male co-housing during the perfuming manipulation.) The reciprocal treatment also showed a significant effect of perfume source: male SC *D. americana* perfumed with *D. novamexicana* males displayed copulation success of 19%, significantly lower success compared to conspecific-perfumed SC *D. americana* (100%, Mann-Whitney U-test: U = 0, *P* = 0.018). For NE-SC pairings, SC *D. americana* male mating rate was also significantly reduced when perfumed with NE *D. americana* males (50%), compared to same-population perfumed SC *D. americana* males (100%, Mann-Whitney U-test: U = 0, *P* = 0.018). In contrast, mating rate of NE *D. americana* males perfumed with SC *D. americana* males increased slightly (56%), but did not significantly differ from the control (same-population-perfumed) males (44%, Mann-Whitney U-test: U = 5.5, *P* = 0.54).

Overall, these results indicate that perfuming males with heteropopulation CHCs influences the frequency of successful copulation with SC *D. americana* females. SC *D. americana* males perfumed with profiles of either *D. novamexicana* or NE *D. americana* had significantly reduced mating rates with their own females. Conversely, perfuming *D. novamexicana* males with SC *D. americana* males significantly increased their mating rate with *D. americana* females, almost completely reversing the pattern of sexual isolation observed for unmanipulated males. This large shift in mating rate for heteropopulation-perfumed *D. novamexicana* indicates that differences in male CHC profiles play a critical role in sexual isolation in the Nov-SC pair. In contrast with the three other classes of heteropopulation-perfumed males, perfuming NE *D. americana* with SC *D. americana* males did not significantly change their mating rate. It possible that this manipulation might have been less effective at altering the CHC profile of NE *D. americana* males (see next section), and/or that mating isolation in the NE-SC pair might depend on more complex factors than a simple shift in CHC composition alone (see Discussion).

### Perfuming shifts CHC composition towards the donor male profile

Using CHCs extracted from an additional set of perfumed males (see Methods), we confirmed that our heteropopulation perfuming manipulation produced quantitative changes in composite male CHC profiles in both the Nov-SC and NE-SC pairs, by shifting male CHCs closer to the donor male profile. PCAs were performed separately for each pairing (N-PCA for the Nov-SC pair, and A-PCA for the NE-SC pair, hereafter), as were analyses of differences among classes of perfumed males. In both cases, of the first three PCs of CHC composition, PC1 (i.e. N-PC1 or A-PC1) varied by both donor and target male identity (Table S7), indicating that perfuming significantly shifted CHC profiles along the primary axis of variation in both perfuming experiments. Nonetheless, the specific donor-target manipulation that was most successful in this regard differed between the Nov-SC and NE-SC pairs. In the Nov-SC pair, *D. novamexicana* males perfumed with SC *D. americana* males significantly differed in CHC composition from control (*D. novamexicana*) perfumed samples (N-PC1) (t(5.9) = -2.8, *P* = 0.031); however, heteropopulation-perfumed SC *D. americana* did not differ from same-population SC *D. americana*-perfumed samples along the same axis (t(3.53) = 1.91, *P* = 0.14). In contrast, in the NE-SC pair, SC *D. americana* perfumed with NE *D. americana* showed a significant shift in A-PC1 (t(4.85) = 4.23, *P* = 0.0088), but there was no significant shift for SC-perfumed NE *D. americana* compared to same-population-perfumed controls (t(4.68) = 0.055, *P* = 0.96). The difference between these pairs might be attributed to high variation seen among samples in some groups (in particular same-population perfumed SC *D. americana*). Regardless, it is clear from these results that this perfuming assay can produce significant, detectable differences in CHC profiles after heteropopulation perfuming, most notably in *D. novamexicana* males.

### Patterns of gene expression and sequence variation implicate an elongase gene that contributes to CHC variation between species

From among a set of 23 candidate genes whose orthologs have functions related to cuticular hydrocarbon variation, both transcriptome and sequence comparisons implicated one specific elongase locus as potentially causal in CHC composition differences between our species. Using whole-transcriptome RNA-seq data from the same three populations (previously generated in Davis and Moyle 2020), we found that 3 of our candidate genes in males and 2 candidates in females were in the upper 10^th^ percentile of genes most differentially expressed between populations (i.e. they showed a stronger species effect than 90% of all 11301 expressed genes in the transcriptome dataset) (Table 3). Of these candidate genes, only CG17821 showed elevated gene expression specifically in SC *D. americana* compared to the other two populations; this pattern was observed in both sexes but is more pronounced in females (Figure 3).

**Figure 3:**
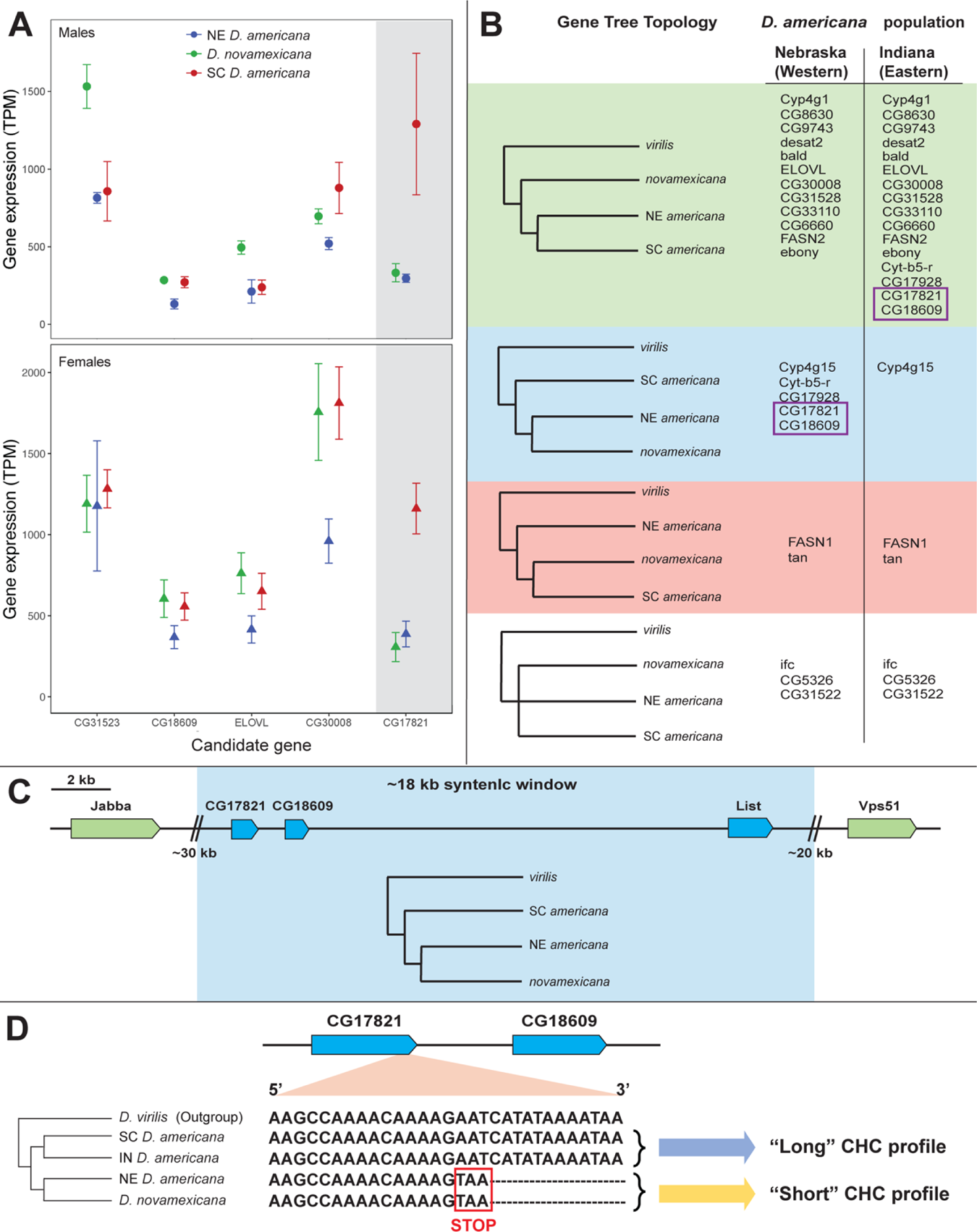
CHC candidate gene expression, genealogical relationships, and sequence variation among *D. americana* group lines. A) Gene expression variation showing mean and standard error between focal species lines for candidate genes with the largest differences in either males (upper, circles) or females (lower, triangles). Gene expression is measured in transcripts per million (TPM, n = 3 for each sample). B) Gene tree topologies for all 23 candidate genes using either western (NE) or eastern (IN) populations for NE *D. americana*. C) Schematic of loci (green/blue boxes) in the genomic window surrounding candidates CG17821 and CG18609. Blue boxes indicate a genealogy that groups the NE *D. americana* population with *D. novamexicana*; gene topology of shaded blue region displayed below line. Green boxes indicate loci with genealogies matching the expected species tree. D) 3’ terminal nucleotides and occurrence of inferred ancestral and derived (truncated) allele in CG17821 among lineages with sequence data.

Closer inspection of our CG17821 sequences revealed that alleles in NE *D. americana* and *D. novamexicana* share a thymine insertion mutation that causes a premature stop codon 4 amino acids upstream of the end of the gene, compared to the annotated gene model in outgroup *D. virilis* and the allele in our SC *D. americana* stock (Figure 3); these four terminal amino acids are not present in RNA transcripts for either NE *D. americana* and *D. novamexicana*. Among our three lines, this suggests an association between the truncated protein product of CG17821 and the “short” CHC phenotype we observe. This association is further supported by additional CHC and sequence data from another *D. americana* population, originally collected in Anderson, IN (species stock center line 15010-0951.00, henceforth IN *D. americana*). Lamb et al. 2020 characterized CHC divergence between females from this IN *D. americana* stock and another *D. novamexicana* line (species stock center 15010-1031.04). The profile for the *D. novamexicana* line had no long compounds (>C30), a low abundance of C29 compounds, and presence of multiple short alkenes (<C27), consistent with the “short” CHC profile we observe for our *D. novamexicana* line. Likewise, the IN *D. americana* profile they observed is broadly consistent with the “long” profile we observed in SC *D. americana*, as both show presence of compounds longer than 30 carbons, a high abundance of C29 length alkenes, and absence of compounds shorter than 27 carbons. In addition, using a genome assembly from the same IN *D. americana* stock (Kim et al. 2020), we found that this population also has the non-truncated allele of CG17821. These data are consistent with our inference that truncation of this allele could contribute to the difference between “long” and “short” CHC phenotypes among populations in this group.

### Evolutionary history of elongase CG17821

Several lines of evidence indicate that CG17821 has a complex evolutionary history within the *D. americana* subgroup, that likely includes post-speciation introgression. The presence of the non-truncated allele in outgroup *D. virilis* (Figure 3) indicates that the truncated allele is derived from a thymine insertion mutation event that took place within the *D. americana* subgroup. Moreover, *D. novamexicana* and our western *D. americana* (NE) line share an identical derived allele at CG17821, indicating an evolutionary history at this locus that disagrees with expected phylogenetic relationships among these three taxa (i.e., where the two *D. americana* populations are expected to be most closely related). Gene trees for each of our 23 candidate genes confirmed that CG17821 has a topology that is discordant with the expected species relationships, placing NE *D. americana* as sister to *D. novamexicana* rather than grouping it with SC *D. americana* (Figure 3). Four other CHC candidate genes also show this discordant topology, most notably CG18609 which is located immediately downstream (within 1kB) of CG17821, according to the annotated genome of *D. virilis* (Figure 3). For 4 out of 5 of these loci—including both CG17821 and CG18609—using the allele from the IN *D. americana* stock instead produces a genealogy where the *D. americana* populations group together, as expected from the species tree (Figure 3).

The observation of phylogenetically discordant sites can be due to several factors, notably incomplete lineage sorting (ILS) or post-speciation introgression (see also Discussion). Two lines of evidence strongly support introgression. First, evidence from SNP variants, across the whole transcriptome and specifically at CG17821, is more consistent with introgression between *D. novamexicana* and our western *D. americana* (NE) population—as determined by Patterson’s D-statistic (Durand et al. 2011). In particular, we found that genome-wide, of 34441 total SNPs detected in 11301 loci within the transcriptome dataset (Davis and Moyle 2020), 14902 supported *D. americana* populations as sister (in accordance with the species tree), 13372 SNPs grouped *D. novamexicana* and NE *D. americana*, and 6167 grouped *D. novamexicana* with SC *D. americana*. The significant excess of shared variants between *D. novamexicana* and our NE *D. americana* line (D = 0.369, P < 0.0001), is consistent with a history of introgression between the progenitors of these two populations. Second, gene tree topologies at and around CG17821 also indicate this specific genomic region shares recent ancestry between *D. novamexicana* and NE *D. americana* due to introgression. CG17821, the adjacent downstream CHC candidate CG18609, and the next nearest gene (List, a neurotransmitter located ∼14 kB downstream) all display shared derived SNPs between *D. novamexicana* and the NE *D. americana* line, and discordant gene tree topologies that group these two lines as sister taxa. In contrast, topologies for the next two nearest up- or down-stream genes in this region reflect the species tree, indicating that recent shared ancestry between *D. novamexicana* and the NE *D. americana* extends across a genomic window between 18 and 68 kB long around the specific region containing CG17821 (Figure 3C). Estimates of linkage disequilibrium (LD) from D. melanogaster autosomes indicate that most LD decays within 200 base pairs, and *r*^2^ (the correlation between SNPs) decreases to <0.1 within a 1 kB window (Franssen et al. 2015); therefore, the size of the putatively-introgressed window observed here is well outside the range expected under ILS, as discordant ancestry due to sorting from ancestral variation is expected in small blocks (Hudson and Coyne 2002, Gao et al. 2015). Note that inversions could also be responsible for capturing ancestrally-segregating variation in larger genomic regions than expected from LD measures, and populations within this group are known to differ in the presence/absence of inversions, including at the specific genomic region containing these genes (Reis et al 2015). However, the inversion at this location is only present in southern *D. americana* (i.e., within the “*D. a. texana*” chromosomal form) to the exclusion of NE *D. americana*, *D. novamexicana*, and *D. virilis*, and therefore could not be responsible for the patterns of discordant variation we observe for these genes here. Coupled with genetic and phenotypic evidence above, these results indicate introgression of these genes between *D. novamexicana* and western *D. americana*—and not ILS—is most likely responsible for their shared allele at CG17821 and therefore potentially for their similarity in “short” CHC phenotypes.

We also evaluated whether there was evidence of elevated protein evolution at CG17821, CG18609 or any other of our candidate genes, based on estimates of the ratio of nonsynonymous to synonymous substitutions (*d*_N_/*d*_S_) at these loci, compared to a genome-wide average rate estimated from our transcriptome-wide dataset (see Methods). For 4397 genes transcriptome-wide, the median and mean *d*_N_/*d*_S_ were 0.065 and 0.0991, respectively (± 0.133 s.d.) (Figure S3). Of our candidate genes, only three had an estimated *d*_N_/*d*_S_ greater than 1 standard deviation above this mean, including the two candidates within the putatively introgressed region—CG17821 (*d*_N_/*d*_S_ = 0.284) and CG18609 (*d*_N_/*d*_S_ = 0.249)—as well as CG6660 (*d*_N_/*d*_S_ = 0.38714)(full results in supplementary Table S8). Notably, these rates of protein change fall within the top 8% of all genes analyzed (Figure S3, all gene data reported in supplementary file). While these estimates are below the threshold for unambiguous evidence of positive selection (*d*_N_/*d*_S_ > 1), only 9 genes in the whole dataset met this criterion. Of the three candidate loci that exhibit elevated protein evolution, CG6660 does not show patterns of genealogical, allelic, or gene expression variation that match our observed CHC phenotypic variation (Figure 3A/B, and above). For both CG17821 and CG18609, we also estimated *d*_N_/*d*_S_ using just *D. virilis* and SC *D. americana* sequence comparisons, to determine if elevated protein evolution in these genes is solely driven by the branch leading to the *D. novamexicana*-NE *D. americana* allele, or if this pattern is more general across the clade. We found our estimate of *d*_N_/*d*_S_ was still modestly elevated between *D. virilis* and SC *D. americana* (*d*_N_/*d*_S_ = 0.284) even when this branch was removed—indicating CG17821 has experienced consistently elevated protein evolution across the whole clade. In contrast, CG18609 protein evolution is estimated to be lower (*d*_N_/*d*_S_ = 0.147) when just considering divergence between *D. virilis* and SC *D. americana*, suggesting that most of the accelerated protein evolution in this gene occurred after the split of the *D. novamexicana*/NE *D. americana* lineage. The timing of elevated protein evolution in CG18609 therefore also generally coincides with CHC profile differences observed here and, like CG17821, CG18609 is also an elongase. However, CG18609 does not show patterns of gene expression that match the “long” versus “short” CHC phenotypes we observe (instead, its expression is significantly reduced in NE *D. americana*; Figure 3A) so while it’s possible that this tandem elongase gene also plays a role in CHC phenotype divergence, it is unlikely to be responsible for the primary axis of CHC variation described here.

Finally, we note that our *d*_N_/*d*_S_ analysis also revealed two odorant binding proteins (OBPs)—to be among the fastest evolving loci in our transcriptome-wide dataset: the orthologs of Obp99d (second fastest in our dataset; *d*_N_/*d*_S_ = 1.589), and *Obp56g* (40^th^ fastest; *d*_N_/*d*_S_ = 0.689) (see supplementary file).

OBPs are thought to facilitate olfactory processing by chaperoning odorants to the olfactory receptor neurons or terminating neuron activity by clearing odorants form the surrounding (Sun et al. 2018). OBP genes have been previously shown to evolve quite rapidly in multiple insect groups (Foret and Maleszka 2006), although rates in these specific OBPs are lower across 6 species in the *Drosophila melanogaster* group (*d*_N_/*d*_S_ > 0.24; Vieira et al. 2007) than we detect here. Notably, both these OBPs are known to expressed in chemosensory organs in D. melanogaster—including adult labellum (*Obp56g*; Galindo and Smith 2001), antenna and maxillary palps (Obp99d; Hekmat-Scafe et al. 2002), and wing sensilla (both loci; He at al. 2019)—consistent with roles in pheromone perception. Moreover, one of these loci—*Obp56g*—is a known seminal fluid protein in the D. melanogaster group (Findlay et al 2008), that also changes in expression within females specifically in response to mating (McGraw et al. 2004). Given these roles, the elevated rates we observe here suggest both genes might contribute to variation in behavioral responses to pheromone and other sexual stimuli among our species (see also discussion).

## Discussion

Identifying genetic and evolutionary mechanisms involved in the earliest steps of reproductive isolation between species is essential for understanding the factors that drive speciation. Evolutionary divergence in sexual signals may be an especially potent contributor to this process (Schemske 2000, Coyne and Orr 2004, Ritchie 2007, Schluter et al. 2009, van Doorn et al. 2009). However, demonstrating the connection between sensory signal divergence and emerging reproductive isolation can be challenging, as it requires identification and demonstration of the direct role of specific signals mediating sexual isolation between species and knowledge of the specific mechanistic changes that have given rise to lineage differences in this signal. Here we have demonstrated that sexual isolation between laboratory populations in the *D. americana* group is based on female choice of male chemical signals, and identified both the specific phenotypic shift between species in pheromone chemistry as well as a genetic variant likely contributing to this phenotypic change and the mating isolation that results from it. Together these data support a clear role for sensory signal divergence in the evolution of premating isolating barriers between populations in this closely related group, and provide insight into how relatively simple genetic and phenotypic mechanisms can cause strong isolation even at early stages of evolutionary divergence.

Female choice of male CHC variation is responsible for isolation between *D. novamexicana* and SC *D. americana*

Differences in copulation success between populations are the product of variation in male choice (via differences in courtship intensity and copulation attempts, depending upon female identity), female choice (via differences in acceptance rates of males, depending upon their identity), or a combination of these factors. Disentangling these alternatives is critical for identifying the specific trait and corresponding preference variation responsible for the emergence of prezygotic species barriers. Here we showed that female choice of a male sensory signal results in strong sexual isolation among *D. americana* group populations, specifically SC *D. americana* female preference for CHC profiles of their own males and rejection of males with alternative profiles. The role of CHCs is exceptionally clear in the case of premating isolation between male *D. novamexicana* and female SC *D. americana*: strong sexual isolation in this cross can be almost entirely reversed by perfuming *D. novamexicana* males with SC *D. americana* male CHCs. In contrast, we find little evidence for differential male preference of females, even though female CHC profiles are also divergent between populations. Instead, our behavioral data indicates that males did not alter their courtship behavior in response to female species identity. Interestingly, these observations also suggest evidence for female choice of male courtship behaviors, whereby SC *D. americana* male courtship consistently induced females to copulate sooner than courtship behaviors of other population males. Moreover, our finding of strong asymmetric mating isolation between male *D. novamexicana* and specific (southeastern) populations of *D. americana* recapitulates patterns of isolation described in this group more than 70 years ago (Spieth 1951, Table S1), indicating that these mating isolation patterns reflect natural variation among populations from which these lines were collected.

These findings fit within a body of studies that have identified either male or female choice of sensory signals as critical for sexual isolation among Drosophila species (patterns reviewed Yukilevich and Patterson 2019). In *Drosophila melanogaster*, many studies have found that male choice of specific female CHC compounds play a role in isolation between closely related heterospecifics (Coyne et al 1995, Billeter et al. 2009) as well as between intraspecific populations (Wu et al. 1995, Hollocher et al. 1997, Yukilevich and True 2008). Such patterns of CHC-mediated male mate-discrimination have also been associated with allelic variation in CHC elongases. For example, D. sechellia reproductive isolation from closely related *D. simulans* results from male mate-discrimination based on female CHC profiles (Shahandeh et al. 2017) and the CHC elongase eloF has been demonstrated to inhibit interspecific mating in the same species (Combs et al. 2018). Female choice of male CHC variation has received comparably less mechanistic attention, however has been demonstrated as a prezygotic barrier in several Drosophila groups, most notably between D. santomea and D. yakuba (Coyne et al. 2002, Mas and Jallon 2005), between the mycophagous D. subquinaria and D. recens (Curtis et al. 2013, Dyer et al. 2013), and between D. mojavensis males reared on different cactus substrates (Havens and Etges 2013). Our results therefore complement a growing body of evidence that shows female choice can be an important factor dictating patterns of sexual isolation among closely related Drosophila populations and species.

### CHC divergence and the role of Elongases

Dissecting the finer details of phenotypic divergence in sensory signals can help pinpoint underlying mechanisms and associated genes responsible, and more clearly demonstrate how these signal changes contribute to isolation between populations. Here we found that CHC divergence between our lines occurs primarily on the basis of compound length, with *D. novamexicana* and NE *D. americana* profiles both having similar enrichment of shorter compounds (“short” phenotype) compared to an enrichment of longer compounds for SC *D. americana* (“long” phenotype). These differences in abundance of shorter-versus longer-chain compounds were observed across both sexes, and across both alkenes and methyl-branched alkanes (Figure 2, Table S6). In contrast, we find no evidence for variation in other features such as double bond or methyl branch location or number. These features themselves suggest that the striking difference between ‘shorter’ and ‘longer’ profile phenotypes could be due to variation in fatty-acid elongase activity, which globally influences the carbon chain length of CHC precursors, and therefore can have consistent downstream effects on both alkenes and methyl-branched alkanes, because both are modified after the elongation step (Pardy et al. 2018).

Strikingly, analyses of gene expression and sequence variation revealed that CG17821 and CG18609—two putative fatty-acid elongases (Szafer-Glusman et al. 2008, Gaudet et al. 2011)—could functionally contribute to our observed variation between longer and shorter CHC profiles. Sequences at both loci are identical between NE *D. americana* and *D. novamexicana*, differ from SC and IN *D. americana* lines, and show modestly elevated protein evolution on the branch leading to *D. novamexicana*/NE *D. americana*, all of which broadly coincide with our observed differences between “short” versus “long” CHC phenotypes. The variation we observed at CG17821 is particularly interesting, because we observe a thymine insertion mutation that results in premature truncation of CG17821 specifically in *D. novamexicana* and NE *D. americana* (Figure 3) and because our gene expression data also indicates reduced expression of the CG17821 allele specifically in these lines. In the latter case, while we might expect gene expression differences to be tissue-specific (that is, observed specifically in the oenocytes: the CHC-producing organs), our data indicate that the population-specific expression signal at this locus is strong enough to detect from whole-body transcriptome data. Both observed sequence and expression changes at CG17821 could individually produce a greater abundance of short CHC products either by lowering elongase protein levels or reducing enzyme activity. Therefore, although our current data cannot differentiate which change may have occurred first (or whether they are pleiotropic), either could produce the pattern of phenotypic difference observed between populations. Overall, these data suggest allelic variation in CG17821 primarily underlies the major axis of CHC divergence observed between SC *D. americana* and the other populations, consistent with a hypothesis of a simple underlying basis for chain length variation. Moreover, because this axis of CHC divergence appears primarily responsible for sexual isolation between *D. novamexicana* and SC *D. americana*, this points to a large role for simple allelic change at this genomic location in the emergence of a strong isolating barrier between these two populations.

### The evolutionary history of CHC divergence and consequences for past and future sexual isolation in this group

Together, our data demonstrate sexual isolation between SC *D. americana* and *D. novamexicana* is due to CHC divergence in compound length and suggest that variation between truncated and non-truncated alleles of the elongase CG17821 contribute to this phenotypic variation. Moreover, both genome-wide SNP variation, as well as localized variation specifically around this locus (Figure 3C), indicate that this phenotypic variation involves a history of introgression. These data point to a model hypothesis for the evolutionary history of transitions involved in the change in CHC profiles between species and, potentially, in the emergence and expression of sexual isolation that depends upon this phenotype.

First, the distribution of both “long” versus “short” CHC phenotypes (shown here and in Lamb et al. 2020), and allelic variation CG17821, indicate that the “short” phenotype and the CG17821 truncated allele are derived states that arose within the *D. americana* group. This shift most likely occurred in western lineages that gave rise to contemporary *D. novamexicana*. The evolutionary forces responsible for the persistence and spread of this phenotype are not yet known. The relationship between variation in environmental factors, insect stress physiology, and features of CHC length and branching is known to be complex. Prior work has associated aspects of CHC divergence with abiotic variation such as latitude (Frentiu and Chenoweth 2010, Rajpurohit et al. 2017, but see Gibbs et al. 2003), and physiological traits such as desiccation resistance (reviewed Chung and Carroll 2015), that suggest a role for natural selection in shaping CHC composition, but the specific sources(s) of selection can be challenging to pinpoint. In the *D. americana* group, species habitats are differentiated primarily on the basis of water availability and these two species, as well as populations within them, differ in key physiological traits such as desiccation resistance (Davis and Moyle 2019). However, sexual selection might also contribute to shaping evolution at CHC loci—as evidenced by the importance of CHC variation for mating success outlined here and elsewhere. We also find evidence that two odorant binding proteins (OBPs)—Obp99d and *Obp56g*—are rapidly evolving across this group. OBPs are known to be important for mediating olfactory behavioral responses during sexual interactions (Laughlin et al. 2008, Leal 2013, Sun et al. 2018) as well as host-plant preference (Matsuo et al. 2007, Comeault et al. 2017). Therefore these loci could be evolving due to natural selection, sexual selection, or both, including in response to changes in pheromone profiles described here, or to other factors such as this group’s close but little investigated habitat association with willow (Salix sp.) trees (Blight and Romano 1953, McAllister 2002, personal observations). Interestingly, our analysis indicates that the elongase CG17821 has experienced modestly elevated protein evolution across the whole *D. virilis* sub-clade; this suggested history of sustained selection indicates this locus might have played important roles in long-term CHC-mediated adaptive divergence across this group. In comparison, the downstream elongase CG18609 has evidence of elevated protein change primarily on the branch leading to *D. novamexicana*/NE *D. americana*, possibly suggesting that this acceleration occurred after impactful changes to CG17821.

Based on our data associating phenotypic divergence with sexual isolation, the appearance of this new CHC phenotype would have reduced sexual compatibility between derived “short” males and females with strong preferences for the “long” ancestral CHC profile. Persistence of this phenotype would have required a broadening of female preference to accommodate males with the derived (“short” CHC) pheromone phenotype (e.g., as has been observed, for example, in male Ostrinia moths; Roelofs et al. 2002). Our data support this expectation, as *D. novamexicana* females are more accepting of both (putatively derived) *D. novamexicana* and (ancestral) SC *D. americana* male CHC phenotypes, while SC *D. americana* females discriminate against derived “short” CHC phenotypes. Interestingly, this model for the evolution of asymmetric sexual isolation is broadly consistent with Kaneshiro’s (1976, 1980) model for peripatric speciation. Kaneshiro observed that females from derived populations frequently have broad preferences for both derived and ancestral male phenotypes; he proposed that this was due to relaxed selection on narrow female preferences in genetically bottlenecked island populations, where founder effects have led to the loss of elements of male courtship. Although Kaneshiro’s model explicitly invokes genetic drift in male trait evolution, our observations indicate his model for the origin of sexual isolation asymmetry could extend more generally to any case where evolutionary change affects a trait important for male sexual signaling. In the case described here, evolutionary change in male CHC profiles (possibly due to selection acting on a CHC elongase gene(s)), accompanied by an apparent broadening of female preferences for these profiles in derived populations, has resulted in the emergence of strong premating asymmetry specifically between females with ancestral trait preferences and males with derived trait values—akin to the model outlined by Kaneshiro.

Intriguingly, our data also support the subsequent movement of this CHC phenotype from *D. novamexicana* into western *D. americana* lineages. One possible explanation for shared variation in the CG17821-CG18609 locus and CHC phenotypes is that this arose from segregating ancestral variation present in both *D. americana* and *D. novamexicana* (that is, is due to ILS). Prior evidence (Caletka and McAllister 2004) as well as data here indicates that some observed site discordance between populations of *D. americana* and *D. novamexicana* is consistent with ILS. However, our analysis strongly supports the additional occurrence of introgression between *D. novamexicana* and western *D. americana*, including specifically of this trait from *D. novamexicana* into western *D. americana* populations. Both their significant excess of shared genome-wide variation (as indicated by the D-statistic), and a genomic region of at least 18 kB of shared recent ancestry surrounding CG17821/CG18609 that accompanies a similar shift to the “short” CHC phenotype in western *D. americana*, support this inference. Note also that while *D. novamexicana* and *D. americana* are reported to be allopatric, limited and sporadic field collections of *D. novamexicana* mean there is an incomplete understanding of the density and extent of this species’ historical range. As a result, this species could have been in closer geographical contact with western *D. americana* populations during the period since their initial split (∼500KYA), which could help explain evidence for introgression after speciation inferred here. Recent work (Sramkowski et al. 2020) describing rare *D. novamexicana*-like pigmentation alleles in geographically disparate *D. americana* populations, similarly suggests evidence of more recent gene exchange between these species.

Given the effect of the “short” CHC phenotype on sexual isolation between *D. novamexicana* and the SC *D. americana* population, this introgression is expected to have consequences for reproductive isolation among *D. americana* lineages. Interestingly, our data indicate that, even though NE *D. americana* shows the general shift to shorter chain CHC length associated with the introgressed region, its current patterns of sexual isolation differ from those seen in *D. novamexicana*. One explanation might be that this introgression-mediated shift in CHCs occurred on a novel *D. americana* genomic background, with potential consequences for the expression of introgressed CHC-affecting loci, and therefore for patterns and strengths of CHC-mediated isolation. A concrete example of these background effects can be seen for CG18609, where NE *D. americana* shares the *D. novamexicana* allele but nonetheless exhibits significantly reduced expression of this locus compared to the other two lineages (Figure 3A). A differential history of allelic exchange (with *D. novamexicana*) across the range of *D. americana*, plus variable genomic background effects on the expression of CHC loci, could contribute to the more complex mating relationships observed here, and elsewhere, among *D. americana* populations. For example, Spieth’s mating analyses (Table S1) identified up to 10-fold variation in mating success among nine disparate populations of *D. americana*. The possibility that this variation is influenced by differential gene exchange with *D. novamexicana* (including of loci with major effects on a signaling phenotype) is testable with work examining mating success between geographically diverse *D. americana* populations—particularly between eastern and western populations—and its covariation with CHC phenotypic and genotypic variation.

Regardless of the collateral consequences for isolation among *D. americana* populations, our data clearly support the role of divergent cuticular hydrocarbon profiles—specifically a general shift in carbon chain length—in sexual isolation between our *D. novamexicana* and SC *D. americana* populations. They also implicate a potentially causal role for the gene CG17821 in determining CHC phenotype via a global change in CHC elongation activity early in the generation of these compounds. These data provide a strong example of how a recently derived allele in a single gene with large phenotypic effects on a sexual signal could underpin asymmetric sexual isolation between closely related species. Moreover, they suggest multiple (behavioral, biochemical, and molecular) lines of evidence that chemosensory processes are evolving rapidly and dynamically across this group.

## Methods

### Experimental Fly stocks

Three stocks were obtained from the University of California San Diego Drosophila Species Stock Center (DSSC): a Drosophila novamexicana stock from San Antonio, NM (15010-1031.08); and two *D. americana* stocks, one from Chadron, NE (15010-0951.06, NE *D. americana* throughout); and one from Jamestown, SC, (15010-1041.29, SC *D. americana* throughout). All stocks were originally collected between 1946 and 1953. *D. americana* has sometimes been divided into two subspecies according to presence (*D. americana* americana) or absence (*D. a. texana*) of a chromosomal fusion of the X- and 4-chromosomes that shows a distinct latitudinal cline (McAllister 2002), however because sub-specific differences apart from this fusion have not been consistently supported, our two lines are treated as populations from within a single heterogeneous species here. The *D. novamexicana* and NE *D. americana* populations used here are the same as those collected and used by Spieth 1951. All fly stocks were reared on standard cornmeal media prepared by the Bloomington Drosophila Stock Center (BDSC) at Indiana University, and were kept at room temperature (∼22 °C). Every assay in this study used virgins isolated within 8 hours of eclosion and aged for 7 days prior to the start of experiments, similar to the 8 days used by Spieth (1951).

### 4x4 unperfumed mating assay

We performed trials in which four virgin males and four virgin females were paired and observed for mating behavior, following the design used by Spieth (1951) which allows for behavioral interactions that might not otherwise be observed in similar single-pair assays. Within each trial, all males are from a single population, as are all females, so are no-choice with respect to the genotype of a mating partner; crosses are varied by pairing males and females of alternative lines. For each trial, 4 males and 4 females were transferred to a single vial without anesthetization and observed for 3 hours.

The number and duration of each copulation event was recorded for each trial. This assay was repeated for a total of 5 replicates for each possible population combination, in reciprocal (i.e., 9 cross types, each of N = 5). Each trial used 7-day old virgins and testing was started within 30 minutes of lights on in the morning.

### Courtship behavior assay

To quantify and evaluate differences in courtship behaviors between cross types, flies were observed in single pair (no-choice) mating assays. We used a modified FlyPi setup—that combines a Raspberry Pi (Raspberry Pi Foundation, Cambridge, UK), pi camera, and 3D printed parts (Chagas et al. 2017)—to record courtship behaviors. Assays were performed in a modified cell culture plate consisting of six 3 cm-diameter culture wells, each with a small amount of cornmeal media in the bottom, that allowed six total crosses to be recorded simultaneously. For each assay, individual virgin male and female flies of a given cross were aspirated without anesthetization to a cell culture well; after 6 total crosses were set up, the plate was videotaped for a contiguous 3-hour period. The six cross combinations assessed in any particular video trial were randomized to account for variance that might otherwise be explained by date. As in the 4x4 mating assay, we performed 5 replicates of all possible population combinations in reciprocal.

Behavioral features were analyzed and scored manually by the same individual to avoid subjective variation among researchers. Three courtship behaviors—male display events, male tapping events, and male licking events—in addition to copulation were scored in each 1x1 trial using the following criteria. A male “display” was counted when a male performed a combination of back and forth movements and occasional wing flicks while maintaining sustained orientation in front of and facing the female. “Tapping” events were defined when a male used his tarsus to touch the abdomen of the female when oriented behind her. “Licking” events were defined when the male’s mouthparts contacted the female genital arch. For male display, tapping, and licking events, individual events were scored separately only if a different behavior (including sitting still/walking away) was observed between instances of the defined behavior. This criterion minimized overcounting of discrete behavioral events, especially those with difficult to view aspects such as number of times a male extruded mouthparts during a contiguous licking event. In addition to these behaviors, copulation success and latency to copulation were also recorded for each trial. Note that females also engage in tapping and other behaviors, but because these appear to be less consistent and are more difficult to observe and score, they were not addressed in this study.

### Extraction and quantification of cuticular hydrocarbons

Cuticular hydrocarbons were extracted from pooled samples by placing five 7-day old virgin flies of a single sex and species identity in a 1.8 mL glass vial (Wheaton 224740 E-C Clear Glass Sample Vials) with 120 μL of hexane (Sigma Aldrich, St Louis, MO, USA) spiked with 10 μg/mL of hexacosane (Sigma Aldrich). After 20 minutes, 100 μL of the solution was removed to a sterilized 1.8 mL glass vial (Wheaton 224740 E-C Clear Glass Sample Vials) and allowed to evaporate overnight under a fume hood. Extracts were stored at -20 °C until analysis. Five replicate samples consisting of 5 flies per sample (25 flies total) were prepared for each sample type, and all replicates were extracted on the same day.

Gas chromatography mass spectrometry (GC/MS) analysis was performed on a 7820A GC system equipped with a 5975 Mass Selective Detector (Agilent Technologies, Inc., Santa Clara, CA, USA) and a HP-5ms column ((5%-Phenyl)-methylpolysiloxane, 30 m length, 250 μm ID, 0.25 μm film thickness; Agilent Technologies, Inc.). Electron ionization (EI) energy was set at 70 eV. One microliter of the sample was injected in splitless mode and analyzed with helium flow at 1 mL/ min. Two different temperature gradients were used depending on the sample type. For CHC analysis of unmanipulated males and females of each species, the following parameters were used: column was set at 50 °C for 0 min, increased to 210 °C at a rate of 35 °C/min, then increased to 280 °C at a rate of 3 °C/min. The MS was set to detect from m/z 33 to 500. For analysis of samples from perfuming trials, the parameters were modified to increase resolution and sensitivity for less abundant compounds: the column was set at 40 °C and held for 3 min, increased to 200 °C at a rate of 35 °C/min, then increased to 280 °C at a rate of 3 °C/min and held for 15 minutes. Chromatograms and spectra were analyzed using MSD ChemStation (Agilent Technologies, Inc.). CHCs were identified on the basis of retention time and electron ionization fragmentation pattern. Compounds are identified in this study as: CXX:Y for alkenes or Me-CXX for methyl-branched alkanes, where XX indicates the length of the carbon chain, and Y indicates number of double bonds, e.g. C21:1 is a 21-carbon alkene with a single double bond.

The abundance of each compound was quantified by normalizing the area under each CHC peak to the area of the hexacosane signal using homebuilt peak selection software (personal correspondence, Dr. Scott Pletcher, Univ. of Michigan).

### Perfuming manipulation and mating assay

‘Perfuming’ involves co-housing target flies in vials filled predominantly with flies of a desired donor identity, so that the CHC profile of the target flies is altered via physical transfer of CHCs from donor flies (Coyne et al. 1994, Dyer et al. 2014, Serrato-Capuchina et al. 2020). Perfuming was performed by placing 2 ‘target’ males with 15 ‘donor’ males within a single vial. All males were 1-day old virgins when perfuming vials were established, and the wings of donor males were removed (under anesthetic) to distinguish them from target males. For the hetero-population perfuming treatments, target males were co-housed with donor males of a different population; same-population (control) treatments paired donor and target males of the same identity. Following 7 days of perfuming, 4 male flies from the same perfuming conditions were transferred (without anesthetization) to vials containing 4 female SC *D. americana*. Within each experiment (Nov-SC, or NE-SC), we ran one trial of each perfuming condition (2 heterospecific-perfumed types, and 2 conspecific-perfumed types) in parallel on the same day (four 4x4 trials in total).

For CHC analysis, males were perfumed as described for the perfumed mating assay, for both Nov-SC and NE-SC experimental pairs, with the exception that 2-3 target males (rather than just 2) were co-housed with 15-18 donor males, enabling 2 parallel perfuming vials to generate 5 target male flies per identity, for each individual CHC extraction. This was then replicated 4 times for each identity to reach N=4 biological replicates for this analysis.

### Candidate gene selection

Our candidate list was generated by searching Flybase (Flybase.org) for annotated *Drosophila melanogaster* genes with one of the protein-coding domains (as identified by InterProt) that have known functions in CHC synthesis (Pardy et al. 2018): “fatty acid desaturase”, “fatty acid desaturase domain”, “Cyp4g”, “ELO family”, or “fatty-acid-synthase”—resulting in an initial list of 34 genes. To this we added the pigmentation genes ebony and tan as they have been shown to alter CHC variation among both D. melanogaster (Massey et al. 2019) and *D. americana* group species (Lamb et al. 2020). With these 36 genes, we identified orthologs in Drosophila virilis (via Flybase using OrthoDB v9.1, Zdobnov et al. 2017) and then evaluated whether each of these loci had transcript expression in our three populations. To do so, we used the BLASTn function of BLAST+ version 2.6.0 (Camacho et al. 2009) to search for matches between the *D. virilis* orthologs and previously published whole-body transcriptome data generated from the same three populations using RNA-seq (Davis and Moyle 2020). Our analyses used only gene expression data from the control (ambient) conditions for both males and females from this study, and excluded any desiccation stress treatment data. Of the 36 initial genes (listed in Table S9), 30 were found to have unambiguous 1-to-1 orthologs in *D. virilis* and, of these, 23 had transcripts present within the Davis and Moyle (2020) dataset. This final set of 23 genes was used to evaluate gene expression and sequence variation among our three lines. For downstream analyses, we also identified alleles of these 23 loci in a genome assembly of *D. americana* Anderson (15010-0951.00, also referred to as A01) generated by Kim et al. 2020, using the BLASTn function of BLAST+ version 2.6.0 (Camacho et al. 2009) with 1/-1 match/mismatch scoring parameters, and retaining the top BLAST hits for each locus.

### Statistical Analyses

All statistical analyses and figure construction in this study was performed with R version 3.4.3. For the unperfumed 4x4 mating dataset, we used Kruskal-Wallis tests to compare copulation success within 3 hours of observation with male or female species identity (one test for each sex). In addition, we used Wilcoxon rank sum tests to perform post-hoc pairwise comparisons for each sex to evaluate which species differed from one another in female acceptance of males or in male courtship success of females. We also calculated the reproductive isolation index RI_1_ for prezygotic barriers as defined by Sobel and Chen (2014). This index is appropriate for comparisons between no-choice tests, and described by the equation RI_1_ = 1 – (heterospecific success)/(conspecific success).

In 1x1 mating trials, mating rate was evaluated using a logistic regression with a chi-square test. For copulation latency, a 2-way ANOVA was performed with latency in minutes as the dependent variable, to test for the effects of male identity, female identity, and their interaction. This was followed by a post-hoc Tukey HSD in which we assessed which male identities (populations) were specifically different for copulation latency. For each of the three courtship behaviors (displays, tappings and lickings), we converted the count data for each trial to a rate per unit time within a trial, by dividing the recorded counts for each trait by the latency to copulation (in minutes) or, if no copulation occurred, the maximum amount of observed time (180 minutes). Describing male behaviors in terms of rates accounts for differences among males in their courtship efficiency; for example, it allows us to differentiate males that performed fewer courtship behaviors because they were rapidly accepted by a female, from males that displayed lower courtship intensity across the total 3 hour monitored period. For each of the three behavior rates (display-rate, tap-rate, lick-rate) as dependent variables, we performed an ANOVA with male identity, female identity, and their interaction as independent variables.

The perfumed 4x4 mating experiment was analyzed using planned contrasts that compared the mating success of males that had the same target identity but different (con-versus hetero-specific) donor perfumes, when paired with SC *D. americana* females. This enabled us to specifically assess the effects of donor perfume variation on mating success of a given target male species. Each pairwise test was performed using a non-parametric Mann-Whitney U test.

Our primary analyses of differences in CHC composition were done after summarizing major axes of variation in CHC phenotypes using a set of Principal Component Analyses (PCA). Separate PCAs were performed on each CHC experiment within this study: one PCA for the initial (unperfumed) parental populations (denoted U-PCA), and one PCA for each of the perfumed experimental pairs—Nov-SC (N-PCA) and NE-SC (A-PCA)(Figure S2). Within each dataset, a one-way ANOVA was used on each of the first 3 PCs to examine effects of sex and species (for the unperfumed dataset) or male perfume identity (in each perfuming study) on CHC composition. Additionally, for the perfumed datasets, T-tests were used to compare differences in PC values between target males that were paired with different (con- and hetero-specific) donor males to assess the effects of our perfuming manipulation on major axes of CHC variation. Factor loadings for N-PC an A-PC analyses, as well as individual compound differences in perfuming pairs can be found in in the supplement (Tables S10, S11, S12 and Figures S1, S2).

For all gene expression analyses, expression is quantified in transcripts per million (TPM), and therefore is normalized within each sample. Here we analyzed datasets separated by sex, as Davis and Moyle (2020) showed that the majority of genes have differential expression based on sex, whereas our primary interest here is in differences between species that might be implicated in female mating choices and male CHC profile variation. With the dataset for each sex, we ran one-way ANOVAs on TPM of every expressed locus (11301 genes total)—including our 23 target candidate genes—to determine loci for which gene expression varied by population. Each gene was ranked according to their resulting F-value (Table 3; un-corrected P-values are given in the supplement (Table S13)). This allowed us to evaluate which candidate genes had more pronounced expression differences between populations, compared to all other genes in the dataset. Candidate genes with greater differential expression between populations than the majority of the transcriptome could suggest these genes impact observed phenotypic differences, even if the number of tests performed make finding significance at alpha = 0.05 difficult.

We also estimated rates of nonsynonymous to synonymous substitutions (*d*_N_/*d*_S_) for 4397 genes in this transcriptome-wide dataset, to quantify molecular evolution transcriptome-wide in this group and to evaluate evidence for positive selection specifically in candidate loci. We used a pipeline for obtaining genome-wide estimates of *d*_N_/*d*_S_ modified from Wu et al. 2018. First, as in Davis and Moyle 2020, transcript sequences from each population were aligned to the *D. virilis* reference genome (Flybase version 1.7, www.flybase.org) to identify loci with expression in the populations studied here. Then, for each gene with transcripts present in all populations and coding sequence (CDS) annotated in the *D. virilis* reference, a consensus fasta was generated for each line (population) for the longest splice variant of a given gene. These consensus fasta sequences were then aligned to each other using PRANK (Löytynoja & Goldman, 2005) with codons enforced and ten bootstrap replicates, allowing us to obtain orthologous gene sequence alignments among the three populations used here and the *D. virilis* outgroup. We then calculated *d*_N_/*d*_S_ from these aligned sequences in PAML v4.9 (Yang 2007) using model M0 in CodeML—a maximum likelihood model for codon substitution. As PAML uses a tree-based model for computing *d*_N_/*d*_S_ that is sensitive to use of the correct gene tree for a given gene (Mendes et al., 2016) consensus gene trees were constructed individually for each gene using RAxML v8.3 with theGTRGAMMA model with 100 bootstraps (Stamatakis, 2014). Lastly, for both CG17821 and CG18609—genes with shared derived alleles for NE *D. americana* and *D. novamexicana*—we also computed the *d*_N_/*d*_S_ value using only sequences for *D. virilis* and SC *D. americana* to determine if observed estimates of protein evolution in these genes is limited to the derived pair or if this pattern is consistent across the clade. Full results of *d*_N_/*d*_S_ for each gene are reported in the supplementary file, with annotation of any known function of orthologs (according to Flybase) for the 100 loci with the highest estimated rates of protein evolution.

To evaluate possible introgression of candidate gene alleles between populations in this group, we constructed additional gene trees for each candidate gene using orthologs from the Anderson IN *D. americana* genome (from Kim et al. 2020) in place of our western (NE) *D. americana* sequences using RAxML v8.3 (Stamatakis, 2014), with alignments created using PRANK. Additional gene trees were generated in the same manner for neighboring genes within 50 kbp around CG17821/CG18609, to estimate size of a window of shared ancestry.

To assess evidence for genome-wide introgression, we used our transcriptome data in conjunction with the *D. virilis* genome to generate a genome-wide set of SNPs for our three focal populations and the *D. virilis* outgroup. To do so, we used bcftools (Li 2011) to generate a VCF (variant calling format) file and call SNP sites between the 4 taxa after filtering for low depth and masking heterozygous sites from individual taxa. With this, we calculated Patterson’s D-statistic (Durand et al. 2011) using the Dtrios program from Dsuite (Malinsky et al. 2020) to calculate the 3 taxa D-statistic as well as an overall P-value using standard errors generated from the default 20 jackknife blocks. We used (((SC americana, NE americana), novamexicana), virilis)(figure 3B top tree) as the expected species tree topology for this analysis.

## Acknowledgements

We would like to thank Matthew Gibson for help with implementing molecular evolution analyses, Zinan Wang for advice about candidate gene selection, Bernard Kim for sequencing the *Drosophila americana* genome and providing assemblies, Scott Pletcher for sharing hydrocarbon analysis software, and Jonathan Massey for connecting JSD and LCM with JYY.

This work was supported by the IU Department of Biology (LCM, JSD), the Department of Defense United States Army Research Office (Grant No. W911NF1610216) and the National Institutes of Health (Grant No. 1P20GM125508) awarded to JYY. The GC/ MS analysis was performed in the UHM Microbial Genetics and Analytical Laboratory (supported by NIH Grant No. 1P20GM125508).

## References

1. Bartelt, R. J., Armold, M. T., Schaner, A. M., & Jackson, L. L. (1986). Comparative Analysis of Cuticular Hydrocarbons in the Drosophila Virilis Species Group. Comparative Biochemistry and Physiology, 83B(4), 731–742.

2. Basolo, A. L. (1995). Phylogenetic evidence for the role of a pre-existing bias in sexual selection. Proceedings. Biological Sciences, 259(1356), 307–311.

3. Baxter, C., Mentlik, J., Shams, I., & Dukas, R. (2018). Mating success in fruit flies: courtship interference versus female choice. Animal Behaviour, 138, 101–108.

4. Billeter, J.C., Atallah, J., Krupp, J. J., Millar, J. G., & Levine, J. D. (2009). Specialized cells tag sexual and species identity in *Drosophila melanogaster*. Nature, 461(7266), 987–991.

5. Blight, W. C., & Romano, A. (1953). Notes on a Breeding Site of *Drosophila americana* Near St . Louis, Missouri. The American Naturalist, 87(833), 111–112.

6. Blomquist G. J., Bagnères A. G. (2010) Insect hydrocarbons: biology. biochemistry, and chemical ecology. Cambridge University Press, New York

7. Bontonou, G., & Wicker-Thomas, C. (2014). Sexual Communication in the Drosophila Genus. Insects, 5(2), 439–458.

8. Butlin, R., Debelle, A., Kerth, C., Snook, R. R., Beukeboom, L. W., Castillo Cajas, R. F., Diao, W., Maan, M. E., Paolucci, S., Weissing, F. J., van de Zande, L., Hoikkala, A., Geuverink, E., Jennings, J., Kankare, M., Knott, K. E., Tyukmaeva, V. I., Zoumadakis, C., Ritchie, M. G., … Schilthuizen, M. (2012). What do we need to know about speciation? Trends in Ecology & Evolution, 27(1), 27–39.

9. Caletka, B. C., & McAllister, B. F. (2004). A genealogical view of chromosomal evolution and species delimitation in the Drosophila virilis species subgroup. Molecular Phylogenetics and Evolution, 33(3), 664–670.

10. Camacho, C., Coulouris, G., Avagyan, V., Ma, N., Papadopoulos, J., Bealer, K., & Madden, T. L. (2009). BLAST+: Architecture and applications. BMC Bioinformatics, 10, 1–9.

11. Chung, H., & Carroll, S. B. (2015). Wax, sex and the origin of species: Dual roles of insect cuticular hydrocarbons in adaptation and mating. BioEssays, 822–830.

12. Chung, H., Loehlin, D., Dufour, H., Vaccarro, K., Millar, J. G., & Carroll, S. B. (2014). A Single Gene Affects Both Ecological Divergence and Mate Choice in Drosophila. Science, 257082.

13. Combs, P. A., Krupp, J. J., Khosla, N. M., Bua, D., Petrov, D. A., Levine, J. D., & Fraser, H. B. (2018). Tissue-Specific cis-Regulatory Divergence Implicates eloF in Inhibiting Interspecies Mating in Drosophila. Current Biology, 28(24), 3969–3975.e3.

14. Coyne, J. A., Kim, S. Y., Chang, A. S., Lachaise, D., & Elwyn, S. (2002). Sexual isolation between two sibling species with overlapping ranges: Drosophila santomea and Drosophila yakuba. Evolution, 56(12), 2424–2434.

15. Coyne, J. A., & Oyama, R. (1995). Localization of pheromonal sexual dimorphism in *Drosophila melanogaster* and its effect on sexual isolation. Proceedings of the National Academy of Sciences of the United States of America, 92(21), 9505–9509.

16. Coyne, J. A., & Orr, H. A., 2004. Speciation. SinauerAssociates, Sunderland, MA

17. Coyne, J. A., Crittenden, A. P., & Maht, K. (1994). Genetics of a Pheromonal Difference Contributing to Reproductive Isolation in Drosophila. Science, 265(5177), 1461–1464.

18. Curtis, S., Sztepanacz, J. L., White, B. E., Dyer, K. a., Rundle, H. D., & Mayer, P. (2013). Epicuticular Compounds of Drosophila subquinaria and D. recens: Identification, Quantification, and Their Role in Female Mate Choice. Journal of Chemical Ecology, 39(5), 579–590.

19. Davis, J. S., & Moyle, L. C. (2020). Constitutive and plastic gene expression variation associated with desiccation resistance differences in the *Drosophila americana* species group. Genes, 11(146).

20. Davis, J. S., & Moyle, L. C. (2019). Desiccation resistance and pigmentation variation reflects bioclimatic differences in the *Drosophila americana* species complex. BMC Evolutionary Biology, 19(1), 1–14.

21. Durand, E. Y., Patterson, N., Reich, D., & Slatkin, M. (2011). Testing for ancient admixture between closely related populations. Molecular Biology and Evolution, 28(8), 2239–2252.

22. Dyer, K. a, White, B. E., Sztepanacz, J. L., Bewick, E. R., & Rundle, H. D. (2014). Reproductive character displacement of epicuticular compounds and their contribution to mate choice in Drosophila subquinaria and Drosophila recens. Evolution; International Journal of Organic Evolution, 68, 1163– 1175.

23. Findlay, G. D., Yi, X., MacCoss, M. J., & Swanson, W. J. (2008). Proteomics reveals novel Drosophila seminal fluid proteins transferred at mating. PLoS Biology, 6(7), 1417–1426.

24. Franssen, S. U., Nolte, V., Tobler, R., & Schlotterer, C. (2015). Patterns of linkage disequilibrium and long range hitchhiking in evolving experimental *Drosophila melanogaster* populations. Molecular Biology and Evolution, 32(2), 495–509.

25. Frentiu, F. D., & Chenoweth, S. F. (2010). Clines in cuticular hydrocarbons in two Drosophila species with independent population histories. Evolution, 64(6), 1784–1794.

26. Gaertner, B. E., Ruedi, E. A., McCoy, L. J., Moore, J. M., Wolfner, M. F., & Mackay, T. F. C. (2015). Heritable variation in courtship patterns in *Drosophila melanogaster*. G3: Genes, Genomes, Genetics, 5(4), 531–539.

27. Gao, Z., Przeworski, M., & Sella, G. (2015). Footprints of ancient-balanced polymorphisms in genetic variation data from closely related species. Evolution, 69(2), 431–446.

28. Gaudet, P., Livstone, M. S., Lewis, S. E., & Thomas, P. D. (2011). Phylogenetic-based propagation of functional annotations within the Gene Ontology consortium. Briefings in Bioinformatics, 12(5), 449–462.

29. Gibbs, A. G., Fukuzato, F., & Matzkin, L. M. (2003). Evolution of water conservation mechanisms in Drosophila. The Journal of Experimental Biology, 206, 1183–1192.

30. Havens, J. A., & Etges, W. J. (2013). Premating isolation is determined by larval rearing substrates in cactophilic Drosophila mojavensis. IX. Host plant and population specific epicuticular hydrocarbon expression influences mate choice and sexual selection. Journal of Evolutionary Biology, 26(3), 562–576.

31. He, Z., Luo, Y., Shang, X., Sun, J. S., & Carlson, J. R. (2019). Chemosensory sensilla of the Drosophila wing express a candidate ionotropic pheromone receptor. PLoS Biology, 17(5), 1–27.

32. Hekmat-Scafe, D. S., Scafe, C. R., McKinney, A. J., & Tanouye, M. A. (2002). Genome-Wide analysis of the odorant-binding protein gene family in *Drosophila melanogaster*. Genome Research, 12(9), 1357– 1369.

33. Hoikkala, A., & Lumme, J. (1987). The Genetic Basis of Evolution of the Male Courtship Sounds in the Drosophila virilis Group. Evolution, 41(4), 827–845.

34. Hollocher, H., Ting, C. T., Wu, M. L., & Wu, C. I. (1997). Incipient speciation by sexual isolation in *Drosophila melanogaster*: Extensive genetic divergence without reinforcement. Genetics, 147(3), 1191–1201.

35. Hudson, R. R., & Coyne, J. A. (2002). Mathematical Consequences of the Genealogical Species. Evolution, 56(8), 1557–1565.

36. Jezovit JA, Levine JD, Schneider J. Phylogeny, environment and sexual communication across the Drosophila genus. J Exp Biol. 2017;220(Pt 1):42–52.

37. Kaneshiro, K. Y. (1980). Sexual Isolation, Speciation and the Direction of Evolution. Evolution, 34(3), 437– 444.

38. Kaneshiro, K. Y. (1976). Ethological Isolation and Phylogeny in the Planitibia Subgroup of Hawaiian Drosophila. Evolution, 30(4), 740.

39. Kim, B. Y., Wang, J. R., Miller, D. E., Barmina, O., Delaney, E., Thompson, A., Comeault, A. A., Peede, D., Agostino, E. R. R. D., Aguilar, J. M., Haji, D., Matsunaga, T., Armstrong, E. E., Zych, M., et. Al. (2020). Highly contiguous assemblies of 101 drosophilid genomes Bernard. BioRxiv.

40. Lamb, A. M., Wang, Z., Simmer, P., Chung, H., Patricia, J., States, U., Lansing, E., States, U., Biology, E., Program, B., Lansing, E., States, U., Biology, E., Program, B., Arbor, A., States, U., & Wittkopp, P. J. (2020). ebony affects pigmentation divergence and cuticular hydrocarbons in *Drosophila americana* and *D. novamexicana*. Frontier in Ecology and Evolution, 8(June), 1–23.

41. Laughlin, J. D., Ha, T. S., Jones, D. N. M., & Smith, D. P. (2008). Activation of Pheromone-Sensitive Neurons Is Mediated by Conformational Activation of Pheromone-Binding Protein. Cell, 133(7), 1255–1265.

42. Leal, W. S. (2013). Odorant reception in insects: Roles of receptors, binding proteins, and degrading enzymes. Annual Review of Entomology, 58, 373–391.

43. Legendre, A., Miao, X. X., Da Lage, J. L., & Wicker-Thomas, C. (2008). Evolution of a desaturase involved in female pheromonal cuticular hydrocarbon biosynthesis and courtship behavior in Drosophila. Insect Biochemistry and Molecular Biology, 38(2), 244–255.

44. Löytynoja, A., & Goldman, N. (2005). An algorithm for progressive multiple alignment of sequences with insertions. Proceedings of the National Academy of Sciences of the United States of America, 102(30), 10557–10562.

45. Li, H. (2011). A statistical framework for SNP calling, mutation discovery, association mapping and population genetical parameter estimation from sequencing data. Bioinformatics, 27(21), 2987– 2993.

46. Malinsky, M., Matschiner, M., & Svardal, H. (2020). Dsuite - fast D-statistics and related admixture evidence from VCF files. BioRxiv, 1–16.

47. Mas, F., & Jallon, J. M. (2005). Sexual isolation and cuticular hydrocarbon differences between Drosophila santomea and Drosophila yakuba. Journal of Chemical Ecology, 31(11), 2747–2752.

48. Massey, J. H., Rice, G. R., Firdaus, A. S., Chen, C. Y., Yeh, S. D., Stern, D. L., & Wittkopp, P. J. (2020). Co-evolving wing spots and mating displays are genetically separable traits in Drosophila. Evolution, 74(6), 1098–1111.

49. Massey, J. H., Akiyama, N., Bien, T., Dreisewerd, K., Wittkopp, P. J., Yew, J. Y., & Takahashi, A. (2019). Pleiotropic Effects of ebony and tan on Pigmentation and Cuticular Hydrocarbon Composition in *Drosophila melanogaster*. Frontiers in Physiology, 10.

50. Matsuo, T., Sugaya, S., Yasukawa, J., Aigaki, T., & Fuyama, Y. (2007). Odorant-binding proteins OBP57d and OBP57e affect taste perception and host-plant preference in Drosophila sechellia. PLoS Biology, 5(5), 0985–0996.

51. McAllister, B. F. (2002). Chromosomal and allelic variation in *Drosophila americana* : selective maintenance of a chromosomal cline. Genome, 45(1), 13–21.

52. McGraw, L. A., Gibson, G., Clark, A. G., & Wolfner, M. F. (2004). Genes Regulated by Mating, Sperm, or Seminal Proteins in Mated Female *Drosophila melanogaster* Lisa. Current Biology, 14, 1509–1514.

53. Morales-Hojas, R., Reis, M., Vieira, C. P., & Vieira, J. (2011). Resolving the phylogenetic relationships and evolutionary history of the Drosophila virilis group using multilocus data. Molecular Phylogenetics and Evolution, 60(2), 249–258.

54. Pardy, J. A., Rundle, H. D., Bernards, M. A., & Moehring, A. J. (2018). The genetic basis of female pheromone differences between *Drosophila melanogaster* and D. simulans. Heredity, 122(1), 93– 109.

55. Patterson, N., Moorjani, P., Luo, Y., Mallick, S., Rohland, N., Zhan, Y., Genschoreck, T., Webster, T., & Reich, D. (2012). Ancient admixture in human history. Genetics, 192(3), 1065–1093.

56. Rajpurohit, S., Hanus, R., Vrkoslav, V., Behrman, E. L., Bergeland, A. O., Petrov, D. A., Cvacka, J., & Schmidt, P. S. (2017). Adaptive dynamics of cuticular hydrocarbon in Drosophila. Journal of Evolutionary Biology, 30(1), 66–80.

57. Reis, M., Vieira, C. P., Lata, R., Posnien, N., & Vieira, J. (2018). Origin and consequences of chromosomal inversions in the virilis group of Drosophila. Genome Biology and Evolution, 10(12), 3152–3166.

58. Ritchie, M. G. (2007). Sexual Selection and Speciation. Annual Review of Ecology, Evolution, and Systematics, 38(1), 79–102.

59. Roelofs, W. L., Liu, W., Hao, G., Jiao, H., Rooney, A. P., & Linn, C. E. (2002). Evolution of moth sex pheromones via ancestral genes. Proceedings of the National Academy of Sciences of the United States of America, 99(21), 13621–13626.

60. Ruedi, E. A., & Hughes, K. A. (2008). Natural genetic variation in complex mating behaviors of male *Drosophila melanogaster*. Behavior Genetics, 38(4), 424–436.

61. Schemske, D. W. (2000). UNDERSTANDING THE ORIGIN OF SPECIES. Evolution, 54(3), 1069–1073.

62. Schluter, D. (2009). Evidence for Ecological Speciation and Its Alternative. Science, 323(5915), 737–741.

63. Seddon, N., Botero, C. A., Tobias, J. A., Dunn, P. O., MacGregor, H. E. A., Rubenstein, D. R., Uy, J. A. C., Weir, J. T., Whittingham, L. A., & Safran, R. J. (2013). Sexual selection accelerates signal evolution during speciation in birds. Proceedings of the Royal Society B: Biological Sciences, 280(1766).

64. Serrato-Capuchina, A., Schwochert, T. D., Zhang, S., & Roy, B. (2020). Pure species discriminate against hybrids in the *Drosophila melanogaster* species subgroup. bioRxiv

65. Shahandeh, M. P., Pischedda, A., & Turner, T. L. (2018). Male mate choice via cuticular hydrocarbon pheromones drives reproductive isolation between Drosophila species. Evolution, 72(1), 123–135.

66. Smadja, C., & Butlin, R. K. (2009). On the scent of speciation: The chemosensory system and its role in premating isolation. Heredity, 102(1), 77–97.

67. Sobel, J. M., & Chen, G. F. (2014). Unification of methods for estimating the strength of reproductive isolation. Evolution, 68(5), 1511–1522.

68. Spieth, H. T. (1951). Mating behavior and sexual isolation in the Drosophila virilis species group. Behaviour, 3(2), 105–145.

69. Sramkoski, L. L., Mclaughlin, W. N., Cooley, A. M., Yuan, D. C., & Wittkopp, P. J. (2020). Genetic architecture of a body color cline in *Drosophila americana*. BioRxiv.

70. Stamatakis, A. (2014). RAxML version 8: A tool for phylogenetic analysis and post-analysis of large phylogenies. Bioinformatics, 30(9), 1312–1313.

71. Sun, J. S., Xiao, S., & Carlson, J. R. (2018). The diverse small proteins called odorant-binding proteins. Open Biology, 8(12).

72. Szafer-Glusman, E., Giansanti, M. G., Nishihama, R., Bolival, B., Pringle, J., Gatti, M., & Fuller, M. T. (2008). A Role for Very-Long-Chain Fatty Acids in Furrow Ingression during Cytokinesis in Drosophila Spermatocytes. Current Biology, 18(18), 1426–1431.

73. Thompson, J. D., Higgins, D. G., & Gibson, T. J. (1994). CLUSTAL W: Improving the sensitivity of progressive multiple sequence alignment through sequence weighting, position-specific gap penalties and weight matrix choice. Nucleic Acids Research, 22(22), 4673–4680.

74. Tomaru, M., Doi, M., Higuchi, H., & Oguma, Y. (2000). Courtship song recognition in the *Drosophila melanogaster* complex: Heterospecific songs make females receptive in D. melanogaster, but not in D. sechellia. Evolution, 54(4), 1286–1294.

75. van Doorn, G. S., Edelaar, P., & Weissing, F. J. (2009). On the Origin of Species by Natural and Sexual Selection. Science, 581(December), 1704–1708.

76. Vieira, F. G., Sánchez-Gracia, A., & Rozas, J. (2007). Comparative genomic analysis of the odorant-binding protein family in 12 Drosophila genomes: Purifying selection and birth-and-death evolution. Genome Biology, 8(11).

77. Wang, Z., Cui, Y., Song, L., & Fang, F. (2016). The hysteresis model of piezoelectric micro-positioning stage based on threshold optimization. PLoS Biology, 38(3), 437–440.

78. Wilkins, M. R., Seddon, N., & Safran, R. J. (2013). Evolutionary divergence in acoustic signals: Causes and consequences. Trends in Ecology and Evolution, 28(3), 156–166.

79. Wu, C. I., Hollocher, H., Begun, D. J., Aquadro, C. F., Xu, Y., & Wu, M. L. (1995). Sexual isolation in *Drosophila melanogaster*: A possible case of incipient speciation. Proceedings of the National Academy of Sciences of the United States of America, 92(7), 2519–2523.

80. Wu, M., Kostyun, J. L., Hahn, M. W., & Moyle, L. C. (2018). Dissecting the basis of novel trait evolution in a radiation with widespread phylogenetic discordance. Molecular Ecology,

81. Yang, Z. (2007). PAML 4: Phylogenetic analysis by maximum likelihood. Molecular Biology and Evolution, 24(8), 1586–1591.

82. Yew, J. Y., & Chung, H. (2017). Drosophila as a holistic model for insect pheromone signaling and processing. Current Opinion in Insect Science, 24, 15–20.

83. Yukilevich, R., & Peterson, E. K. (2019). The evolution of male and female mating preferences in Drosophila speciation. Evolution, 73(9), 1759–1773.

84. Yukilevich, R., & True, J. R. (2008). Incipient sexual isolation among cosmopolitan *Drosophila melanogaster* populations. Evolution, 62(8), 2112–2121.

85. Zdobnov, E. M., Tegenfeldt, F., Kuznetsov, D., Waterhouse, R. M., Simao, F. A., Ioannidis, P., Seppey, M., Loetscher, A., & Kriventseva, E. V. (2017). OrthoDB v9.1: Cataloging evolutionary and functional annotations for animal, fungal, plant, archaeal, bacterial and viral orthologs. Nucleic Acids Research, 45(D1), D744–D749.

