## Supplementary material for "A shift to shorter cuticular hydrocarbons accompanies sexual isolation among *Drosophila americana* group populations"

**Table S1: Mating rate data from Table 3 of Spieth 1951. Values are proportion of females mated from 4 male x 4 female mating trials (N=4 replicates per cross). Intra-population crosses are shaded along the diagonal. Spieth used three designations for different populations of *D. americana*, including the subspecific names. *D. novamexicana* Sa and *D. a. americana* Cd are the same populations as the ones used in this study (*D. novamexicana* and NE *D. americana,* respectively, in our study), and are bolded. SC *D. americana* was not used by Spieth, but shares geographic and chromosomal similarity to the No and Lk *D. a. texana* populations reported here.**

|  | Males | *D. novamexicana* | | *D. a. americana* (Western) | | | | *D. a. americana* (Eastern) | | | *D. a. texana* | |
| --- | --- | --- | --- | --- | --- | --- | --- | --- | --- | --- | --- | --- |
| Females |  | **Sa** | Wh | Ch 1 | Ch 2 | Po | **Cd** | Id | Sm | Mi | No | Lk |
| *D. novamexicana* | **Sa** | **0.80** | 0.40 |  |  | 0.562 | **0.375** |  | 0.067 | 0.687 | **0.865** | **0.50** |
|  | Wh | 0.266 | 0.625 |  |  | 0.066 | 0.125 |  | 0.133 | 0.437 | 0.563 | 0.467 |
| *D. a. americana* (Western) | Ch1 |  |  | 0.643 | 0.50 | 0.812 | 0.715 | 0.308 | 0.936 | 0.663 | 0.865 |  |
|  | Ch2 |  |  | 0.466 | 0.715 | 0.60 | 0.461 | 0.80 | 0.833 | 0.537 | 1.00 |  |
|  | Po | 0.312 | 0.302 | 0.357 | 0.60 | 0.857 | 0.50 | 0.536 | 0.20 | 0.532 | 0.676 | 0.50 |
|  | **Cd** | **0.438** | 0.143 | 0.133 | 0.45 | 0.50 | **0.436** | 0.375 | 0.375 | 0.60 | 0.57 | 0.375 |
| *D. a. americana* (Eastern) | Id |  |  | 0.00 | 0.133 | 0.312 | 0.133 | 0.625 | 0.188 | 0.25 | 0.40 |  |
|  | Sm | 0.125 | 0.00 | 0.063 | 0.00 | 0.00 | 0.25 | 0.125 | 0.374 | 0.467 | 0.40 | 0.063 |
|  | Mi | 0.067 | 0.063 | 0.23 | 0.50 | 0.466 | 0.50 | 0.438 | 0.374 | 0.50 | 0.734 | 0.357 |
| *D. a. texana* | No | **0.00** | 0.67 | 0.125 | 0.063 | 0.066 | 0.267 | 0.600 | 0.187 | 0.932 | 0.75 | 0.563 |
|  | Lk | **0.125** | 0.00 |  |  | 0.937 | 0.562 |  | 0.533 | 0.688 | 0.688 | 0.875 |

**Table S2: Posthoc comparisons for differences in male courtship success (based on proportion mated) between male populations regardless of female identity. Pairwise Wilcoxon rank sum tests were performed for each pair with Benjamini-Hochberg corrected *P*-values reported.**

| **Male pop. pair** | ***P-adj*** |
| --- | --- |
| **Nov-NE** | 0.88 |
| **Nov-SC** | 0.73 |
| **NE-SC** | 0.73 |

**Table S3: Posthoc comparisons of pairwise species differences in male behavior rates. Tukey HSD post-hoc mean differences and *P*-values between male identities for each behavior.**

| **Pop. Pair** | **Nov-NE** | | **Nov-SC** | | **NE-SC** | |
| --- | --- | --- | --- | --- | --- | --- |
|  | **mean diff.** | ***P*** | **mean diff.** | ***P*** | **mean diff.** | ***P*** |
| **Display-rate** | -0.0088 | 0.93 | 0.061 | 0.037 | 0.052 | 0.086 |
| **Tap-rate** | -0.0076 | 0.93 | 0.061 | 0.023 | 0.053 | 0.052 |
| **Lick-rate** | -0.0059 | 0.99 | 0.10 | 0.11 | 0.094 | 0.14 |

**Table S4: Posthoc comparisons of pairwise species differences for copulation latency based on male population identity. Tukey HSD post-hoc mean differences and *P*-values between male identities reported.**

| **Male pop. pairs** | **mean diff.** | ***P-adj*** |
| --- | --- | --- |
| **Nov-NE** | 15.93 | 0.81 |
| **Nov-SC** | -42.33 | 0.07 |
| **NE-SC** | -58.28 | 0.23 |

**Table S5: Loadings for principal component (U-PC) axes for each individual compound detected from GC/MS analysis of unmanipulated (unperfumed) samples.**

|  | **PC1** | **PC2** | **PC3** | **PC4** | **PC5** |
| --- | --- | --- | --- | --- | --- |
| **C21:1** | 0.038367 | -0.60597 | -0.4935 | -0.20113 | 0.438403 |
| **C23:1** | 0.333165 | 0.0426 | 0.21122 | 0.086868 | 0.125511 |
| **C25:1** | 0.325075 | 0.061267 | 0.31656 | -0.01855 | 0.320399 |
| **C27:1** | 0.239535 | -0.40331 | 0.407701 | -0.30115 | -0.47207 |
| **C27:3** | 0.32221 | -0.13909 | 0.167199 | -0.27308 | 0.14094 |
| **C29:1** | -0.32568 | -0.14604 | 0.131973 | -0.05233 | -0.03849 |
| **C29:3** | -0.31563 | -0.1888 | -0.02265 | -0.1801 | -0.47663 |
| **C31:2** | -0.32453 | -0.13391 | -0.00704 | -0.077 | 0.178389 |
| **Me-C24** | 0.307398 | -0.07295 | -0.33943 | 0.631199 | -0.32662 |
| **Me-C26** | 0.279074 | -0.35624 | 0.090653 | 0.162503 | -0.01055 |
| **Me-C28** | -0.19882 | -0.48769 | 0.227287 | 0.465292 | 0.051521 |
| **Me-C30** | -0.31265 | -0.01546 | 0.473697 | 0.322454 | 0.277043 |
| Notation: CXX:Y for alkenes or Me-CXX for methyl-branched alkanes, where XX indicates the length of the carbon chain, and Y indicates number of double bonds. | | | | | |

**Table S6: Analyses of individual cuticular hydrocarbon compounds in unmanipulated (unperfumed) samples, from two-way ANOVAs testing for population or sex differences. Compound abundance values were log-transformed for analysis.**

|  | **population** | | **sex** | |
| --- | --- | --- | --- | --- |
|  | ***F*-value** | ***P-*value** | ***F*-value** | ***P-*value** |
| **C21:1**† | 6.96 | 0.0098 |  |  |
| **C23:1** | 141.84 | < 0.0001* | 1.31 | 0.26 |
| **C25:1** | 59.59 | < 0.0001* | 7.60 | 0.011 |
| **C27:1** | 20.87 | < 0.0001* | 36.83 | < 0.0001* |
| **C27:3** | 103.34 | < 0.0001* | 33.71 | < 0.0001* |
| **C29:1** | 716.92 | < 0.0001* | 0.12 | 0.73 |
| **C29:3** | 214.21 | < 0.0001* | 0.036 | 0.85 |
| **C31:2** | 49.41 | < 0.0001* | 1.76 | 0.20 |
| **Me-C24** | 39.38 | < 0.0001* | 14.50 | < 0.0001* |
| **Me-C26** | 55.85 | < 0.0001* | 84.72 | < 0.0001* |
| **Me-C28** | 25.00 | < 0.0001* | 22.51 | < 0.0001* |
| **Me-C30** | 156.21 | < 0.0001* | 11.73 | 0.0020* |
| *Bonferroni-corrected significance level is *P* < 0.0042  †Only males have C21:1, therefore this ANOVA included a species effect only. | | | | |

**Table S7: Analyses of the first three PCs of CHC compound variation from each perfuming pair, from one-way ANOVAs examining the effects of donor and target male identity on each PC.**

| **Male** | **Donor** | | **Target** | |
| --- | --- | --- | --- | --- |
| **Nov-SC pair** | ***F*-value** | ***P*-value** | ***F*-value** | ***P*-value** |
| N-PC1 | 10.71 | **0.0060** | 16.21 | **0.0015** |
| N-PC2 | 0.91 | 0.36 | 1.21 | 0.29 |
| N-PC3 | 0.009 | 0.93 | 0.003 | 0.96 |
| **NE-SC pair** |  |  |  |  |
| A-PC1 | 7.49 | **0.017** | 18.40 | **0.00088** |
| A-PC2 | 0.065 | 0.80 | 1.75 | 0.21 |
| A-PC3 | 1.38 | 0.27 | 1.0 | 0.36 |

**Table S8: Clade-wide *d*_N_/*d*_S_ values for 23 candidate genes associated with CHC function. For genes with elevated clade-wide *d*_N_/*d*_S_, we evaluated *d*_N_/*d*_S_ between *D. virilis* and SC *D. americana* only to determine if accelerated evolution is limited to the most derived pairs.**

| **Candidate gene** | **motif** | ***d*_N_/*d*_S_** | ***d*_N_/*d*_S_ *D. virilis* – SC *D. americana*** |
| --- | --- | --- | --- |
| Cyp4g1 | Cyp4g | 0.0469 |  |
| Cyp4g15 | Cyp4g | 0.0862 |  |
| Cyt-b5-r | desat | 0.0001 |  |
| ifc | desat | 0.0266 |  |
| CG17928 | desat | 0.133 |  |
| CG8630 | desat | 0.118 |  |
| CG9743 | desat | 0.0712 |  |
| desat2 | desat | 0.0780 |  |
| Baldspot | elongase | 0.0280 |  |
| bond | elongase | 0.0503 |  |
| ELOVL | elongase | 0.0191 |  |
| CG5326 | elongase | 0.0081 |  |
| **CG17821** | **elongase** | **0.284** | **0.28423** |
| **CG18609** | **elongase** | **0.249** | **0.14718** |
| CG30008 | elongase | 0.151 |  |
| CG31522 | elongase | 0.0422 |  |
| CG31523 | elongase | 0.0410 |  |
| CG33110 | elongase | 0.0178 |  |
| **CG6660** | **elongase** | **0.387** |  |
| FASN1 | fatty-acid-synthase | 0.0274 |  |
| FASN2 | fatty-acid-synthase | 0.0979 |  |
| ebony | pigment | 0.0280 |  |
| tan | pigment | 0.0269 |  |
| Note: **Bold** genes indicate *d*_N_/*d*_S_ values over 1 standard deviation from mean *d*_N_/*d*_S_ value of 4396 genes analyzed. | | |  |

**Table S9: List of candidate genes and presence/absence of transcript expression in transcriptomes of each population used here (as determined by BLASTn hits to *D. virilis* ortholog).**

| **Candidate gene** | ***D. virilis* ortholog** | **InterProt domain** | ***D. novamexicana*** | **SC *D. americana*** | **NE *D. americana*** |
| --- | --- | --- | --- | --- | --- |
| Cyp4g1 | GJ15981 | Cyp4g | X | X | X |
| Cyp4g15 | GJ19152 | Cyp4g | X | X | X |
| Cyt-b5-r | Cyt-b5-r | fatty-acid desaturase | X | X | X |
| ifc | GJ15437 | fatty-acid desaturase | X | X | X |
| CG15531 | GJ10409 | fatty-acid desaturase |  |  |  |
| CG17928 | GJ17306 | fatty-acid desaturase | X | X | X |
| CG8630 | GJ10413 | fatty-acid desaturase | X | **X** | X |
| CG9743 | GJ10408 | fatty-acid desaturase | X | X | X |
| CG9747 | GJ10410 | fatty-acid desaturase |  |  |  |
| desat1 | desat1 | fatty-acid desaturase |  |  |  |
| desat2 | desat2 | fatty-acid desaturase | X | X | X |
| Baldspot | GJ12451 | elongase | X | X | X |
| bond | GJ24166 | elongase | X | X | X |
| **ELOVL** | **GJ24115** | **elongase** | **X** | X | X |
| Elo68β | GJ12287 | elongase |  |  |  |
| CG5326 | GJ24664 | elongase | X | X | X |
| **CG17821** | **GJ22070** | **elongase** | X | **X** | **X** |
| **CG18609** | **GJ22071** | **elongase** | X | **X** | X |
| **CG30008** | **GJ22296** | **elongase** | **X** | X | **X** |
| CG31522 | GJ24118 | elongase | X | X | X |
| **CG31523** | **GJ24117** | **elongase** | X | **X** | X |
| CG33110 | GJ24167 | elongase | X | X | X |
| **CG6660** | **GJ14350** | **elongase** | X | X | X |
| sit | GJ23058 | elongase |  |  |  |
| FASN1 | GJ21736 | fatty-acid-synthase | X | X | X |
| FASN2 | GJ21725 | fatty-acid-synthase | X | X | X |
| FASN3 | GJ22868 | fatty-acid-synthase |  |  |  |
| ebony | GJ14444 | pigment | X | X | X |
| tan | GJ14875 | pigment | x | X | X |

**Table S10: Nov-SC perfume sample loadings for principal component (N-PC) axes for each individual compound detected from GC/MS analysis.**

| **Compound** | **PC1** | **PC2** | **PC3** | **PC4** | **PC5** |
| --- | --- | --- | --- | --- | --- |
| **C21:1** | -0.19779 | 0.404459 | -0.3266 | 0.137685 | 0.558152 |
| **C23:1** | -0.30656 | -0.05955 | -0.00618 | -0.34054 | 0.070847 |
| **9-C25:1** | -0.30779 | -0.00321 | -0.04702 | -0.2045 | -0.32963 |
| **7-C25:1** | -0.31142 | 0.06597 | 0.030394 | -0.09894 | 0.101648 |
| **C26:1** | -0.29282 | 0.204916 | -0.15414 | -0.00641 | -0.10933 |
| **9-C27:1** | -0.27277 | 0.298372 | -0.02433 | 0.048271 | 0.027908 |
| **7-C27:1** | -0.20702 | 0.376598 | 0.433784 | 0.573061 | -0.10863 |
| **C27:3** | -0.30556 | 0.105753 | -0.06235 | -0.19079 | -0.17517 |
| **C29:1** | 0.252922 | 0.291222 | -0.23988 | -0.3893 | 0.298844 |
| **C29:3** | 0.206667 | 0.360967 | -0.57894 | 0.13495 | -0.52396 |
| **Me-C24** | -0.28106 | -0.19395 | -0.11763 | -0.14644 | -0.29043 |
| **Me-C26** | -0.30827 | 0.101195 | 0.131457 | -0.14341 | 0.0423 |
| **Me-C28** | 0.188634 | 0.411308 | 0.461127 | -0.47413 | -0.09536 |
| **Me-C30** | 0.250042 | 0.335169 | 0.193061 | -0.09859 | -0.22396 |

**Table S11: NE-SC perfume sample loadings for principal component (A-PC) axes for each individual compound detected from GC/MS analysis.**

|  | **PC1** | **PC2** | **PC3** | **PC4** | **PC5** |
| --- | --- | --- | --- | --- | --- |
| **C21:1** | -0.09441 | 0.00287 | 0.794305 | -0.11209 | -0.05871 |
| **C23:1** | -0.30922 | 0.055218 | 0.099506 | 0.146023 | -0.36424 |
| **9-C25:1** | -0.31683 | 0.079214 | 0.005997 | 0.078611 | -0.16968 |
| **7-C25:1** | -0.31617 | 0.07004 | 0.055269 | 0.112009 | -0.21679 |
| **C26:1** | -0.30023 | 0.221801 | 0.10154 | -0.08195 | 0.108149 |
| **9-C27:1** | -0.25244 | 0.400166 | -0.03123 | -0.2169 | 0.377844 |
| **7-C27:1** | 0.032287 | 0.583285 | -0.34794 | 0.430135 | -0.17495 |
| **C27:3** | -0.29941 | 0.215362 | -0.05762 | -0.21199 | 0.212106 |
| **C29:1** | 0.252804 | 0.291122 | 0.345164 | 0.213874 | -0.10676 |
| **C29:3** | 0.227292 | 0.305015 | -0.09092 | -0.69252 | -0.59159 |
| **Me-C24** | -0.3067 | -0.08117 | 0.087312 | 0.258352 | -0.34636 |
| **Me-C26** | -0.30604 | 0.168255 | 0.031321 | -0.16821 | 0.199741 |
| **Me-C28** | 0.267628 | 0.294868 | 0.204858 | 0.200103 | 0.113134 |
| **Me-C30** | 0.27504 | 0.300333 | 0.206013 | 0.057128 | 0.154919 |

**Table S12: Analyses of individual cuticular hydrocarbon compounds in perfumed samples, from t-tests comparing target males with con- and hetero-specific donor perfumes. For example, *D. novamexicana* C21:1 tests the difference between C21:1 values in *D. novamexicana*** **target** **males perfumed with *D. novamexicana* or SC *D. americana* donor males. Compound abundance values were log-transformed for analysis.**

| **Experiment** | **Nov-SC** | | | | **NE-SC** | | | |
| --- | --- | --- | --- | --- | --- | --- | --- | --- |
| **Target** | ***D. novamexicana*** | | **SC *D. americana*** | | **NE *D. americana*** | | **SC *D. americana*** | |
|  | **t-score** | ***P*-value** | **t-score** | ***P*-value** | **t-score** | ***P*-value** | **t-score** | ***P*-value** |
| **C21:1** | -1.82 | 0.14 | 0.089 | 0.93 | -3.97 | **0.027** | 2.19 | 0.079 |
| **C23:1** | 3.74 | **0.014** | 3.35 | **0.037** | -1.65 | 0.15 | 6.79 | **0.0064** |
| **9-C25:1** | 5.05 | **0.0053** | 4.82 | **0.0053** | 0.80 | 0.46 | 5.70 | **0.0091** |
| **7-C25:1** | 4.23 | **0.0061** | 2.62 | **0.044** | -2.18 | 0.11 | 4.81 | **0.015** |
| **C26:1** | 1.15 | 0.29 | 1.62 | 0.17 | -0.89 | 0.42 | 3.83 | **0.013** |
| **9-C27:1** | 1.54 | 0.17 | 1.18 | 0.32 | 0.56 | 0.60 | 3.58 | **0.018** |
| **7-C27:1** | 0.85 | 0.45 | 0.013 | 0.99 | 0.077 | 0.94 | 1.44 | 0.21 |
| **C27:3** | 4.55 | **0.0039** | 4.07 | **0.011** | 1.89 | 0.14 | 4.76 | **0.0060** |
| **C29:1** | -12.76 | **0.00077*** | -1.42 | 0.22 | -14.27 | **0.00077*** | -1.25 | 0.28 |
| **C29:3** | -1.73 | 0.18 | -0.26 | 0.81 | 0.34 | 0.75 | 0.90 | 0.41 |
| **Me-C24** | 3.68 | **0.024** | 2.28 | 0.073 | -1.27 | 0.26 | 4.63 | **0.018** |
| **Me-C26** | 3.91 | **0.0079** | 2.21 | 0.11 | 0.89 | 0.41 | 4.29 | **0.0090** |
| **Me-C28** | -2.29 | 0.074 | -0.85 | 0.44 | -3.47 | **0.017** | -0.37 | 0.73 |
| **Me-C30** | -5.57 | **0.0044** | -1.98 | 0.098 | -4.16 | **0.0059** | -1.52 | 0.18 |
| * Reached Bonferroni-corrected significance level of *P* < 0.0036 **Bold** indicates significance level *P* < 0.05 | | | | | | | | |

**Table S13: Species differences in quantitative gene expression for 23 candidate genes associated with CHC function, analyzed separately by sex, with *F*-value and un-adjusted *P*-value of ANOVA test.**

|  |  | Males | Females |
| --- | --- | --- | --- |

| **Candidate gene** | ***D. virilis* ortholog** | ***F*-value*** | ***P-*value** | ***F*-value*** | ***P-*value** |
| --- | --- | --- | --- | --- | --- |
| Cyp4g1 | GJ15981 | 0.143 | 0.87 | 0.158 | 0.86 |
| Cyp4g15 | GJ19152 | 2.07 | 0.22 | 1.05 | 0.41 |
| Cyt-b5-r | Cyt-b5-r | 2.32 | 0.19 | 0.736 | 0.51 |
| ifc | GJ15437 | 0.122 | 0.89 | 0.74 | 0.52 |
| CG17928 | GJ17306 | 1.09 | 0.40 | 0.0233 | 0.98 |
| CG8630 | GJ10413 | 3.48 | 0.11 | 1.46 | 0.30 |
| CG9743 | GJ10408 | 2.83 | 0.15 | 1.75 | 0.25 |
| desat2 | desat2 | 0.216 | 0.81 | 0.903 | 0.45 |
| Baldspot | GJ12451 | 0.814 | 0.49 | 0.916 | 0.45 |
| bond | GJ24166 | 0.710 | 0.38 | 0.949 | 0.76 |
| **ELOVL** | **GJ24115** | **9.61** | 0.019 | 2.67 | 0.15 |
| CG5326 | GJ24664 | 4.14 | 0.087 | 0.138 | 0.87 |
| **CG17821** | **GJ22070** | 3.50 | 0.11 | **17.2** | 0.0033 |
| **CG18609** | **GJ22071** | **8.19** | 0.026 | 1.85 | 0.24 |
| **CG30008** | **GJ22296** | 2.19 | 0.21 | **4.33** | 0.069 |
| CG31522 | GJ24118 | 0.744 | 0.52 | 0.380 | 0.70 |
| **CG31523** | **GJ24117** | **6.57** | 0.040 | 0.0481 | 0.95 |
| CG33110 | GJ24167 | 0.466 | 0.65 | 0.436 | 0.67 |
| **CG6660** | **GJ14350** | 1.15 | 0.39 | 1.20 | 0.36 |
| FASN1 | GJ21736 | 1.42 | 0.32 | 0.526 | 0.62 |
| FASN2 | GJ21725 | 1.83 | 0.25 | 1.02 | 0.41 |
| ebony | GJ14444 | 0.673 | 0.22 | 1.15 | 0.38 |
| tan | GJ14875 | 1.49 | 0.31 | 0.347 | 0.72 |
| *Note: **Bold** genes in these columns indicate genes in the top 10% of differentially expressed genes within sex. | | | | | |

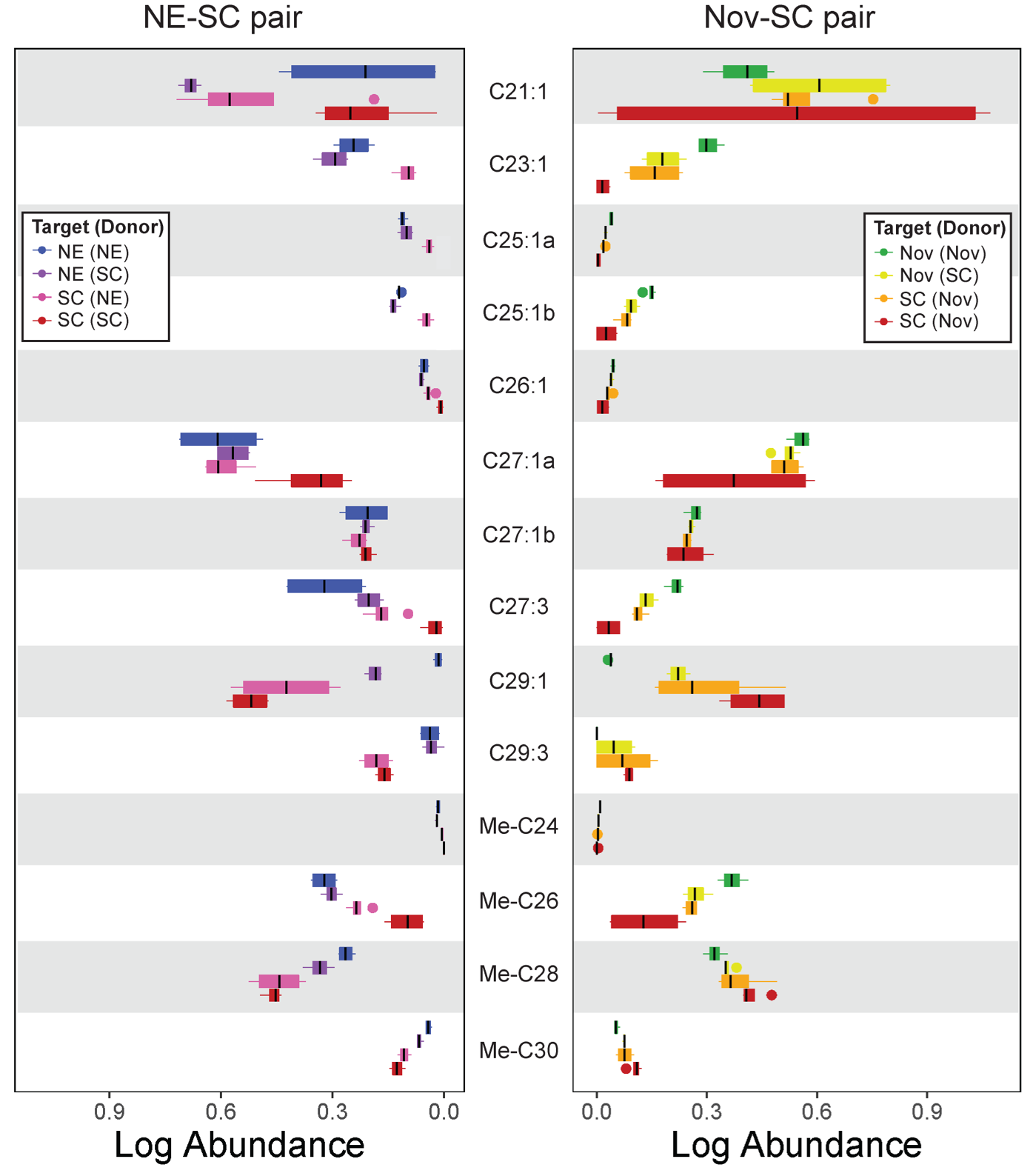

**Figure S1: Relative log-scale abundance of compounds for perfumed males from NE-SC (left) and Nov-SC (right) experimental pairs for each major compound type (line = mean, box = central quartile, whiskers = S.E, *n* = 4 for all bars). Legends indicate perfuming identities within each experiment, with the target male listed first, and the donor male in parentheses. A total of 10 alkenes were detected, including 4 that were not detected in the unperfumed analysis; however, very long chain C31 molecules (found only in SC *D. americana*; Figure 3) were absent in these analyses. Note that SC *D. americana* has the same conspecific (control) perfume treatment in both experiments, but these control samples were gathered and analyzed separately in each. Tests for species differences for individual compounds are shown in Table S7.**

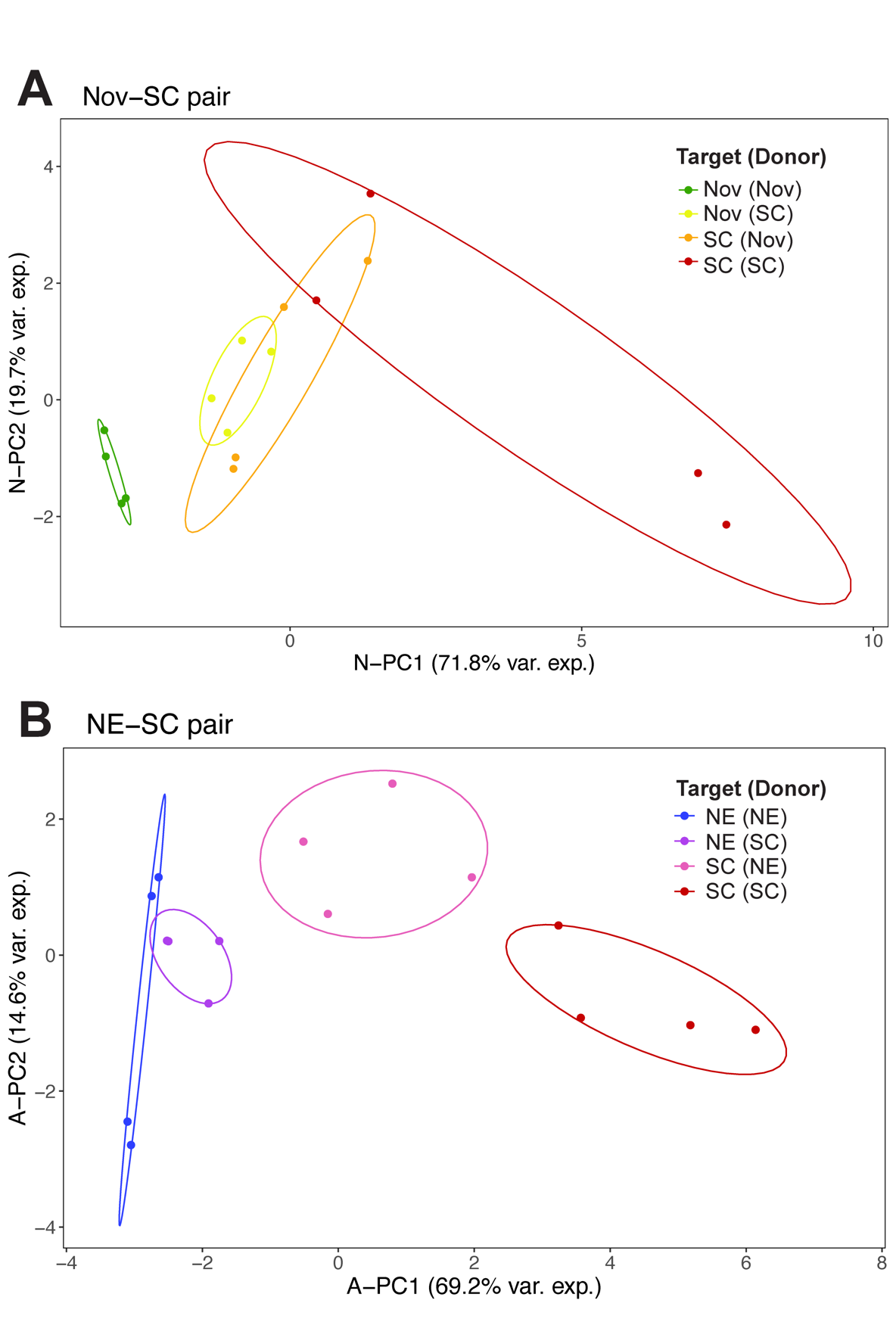

**Figure S2: PC1 and PC2 axes showing individual samples for each male perfuming identity for the Nov-SC pair (panel A) and the NE-SC pair (panel B). Legends indicate perfuming identities within each experiment, with the target male listed first, and the donor male in parentheses. Nov-SC perfuming significantly shifted N-PC1 value for *D. novamexicana* flies (green-yellow pair) but not for perfumed SC *D. americana* (orange-red pair). Conversely, NE-SC perfuming significantly shifted the A-PC1 axis (red-magenta pair) but not for perfumed *D. americana* males (blue-purple pair). Ellipses indicate 90% bivariate normal density for each perfume identity.**

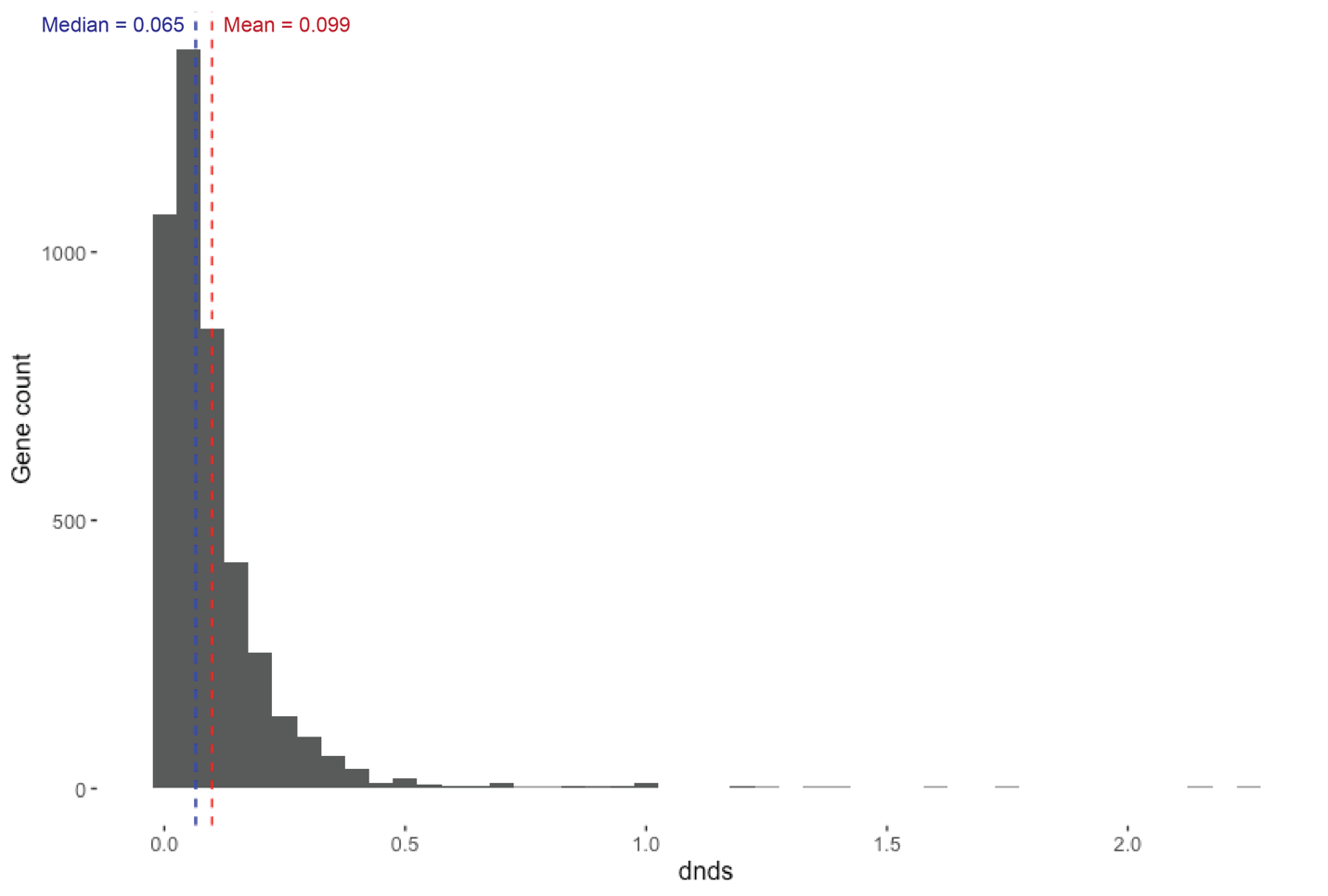

**Figure S3: Distribution of clade-wide *d*_N_/*d*_S_ values from 4397 genes. The median *d*_N_/*d*_S_ value was 0.065, while the mean and standard deviation were 0.0991 ± 0.133.**
